## Supplementary Material for "Combined physical and pharmacological anabolic osteoporosis therapies increase bone response and mechanoregulation in female mice"

**Table S.1**: Color encoding for significant differences to a specific group. Significances are depicted with a symbol in the color of the corresponding group using two-way ANOVA for the shown groups followed by Tukey’s post-hoc test (p<0.05: **o**: to monotreatment, ●: to combined treatment, ○○○○○●●●: to VEH 0N, to VEH 8N, to BIS 0N, to PTH 0N, to ASC 0N, to BIS 8N, to PTH 8N, to ASC 8N).

| ○ | ○ | ○ | ○ | ○ | ● | ● | ● |
|---|---|---|---|---|---|---|---|
| VEH 0N | VEH 8N | BIS 0N | PTH 0N | ScIAB 0N | BIS 8N | PTH 8N | ScIAB 8N |

**Table S.2**: Static morphometric and mechanical parameters of the trabecular compartment of non-loaded (0N) and loaded (8N) 6th caudal vertebrae at weeks 15, 20, 22, and 24 (w15, w20, w22, w24). Values are presented as group means ± standard error, Δ represents the %-change from week 20 to 24 (Δw20-24). Two-way ANOVA over time for the shown groups followed by Tukey’s post-hoc test was used on Δw20-24. ‘x’ denotes the reference group; asterisks (*, **, ***) indicate groups that are significantly different from 'x'. Significance is indicated as: *p<0.05, **p<0.01, ***p<0.001. Synergy (S) or antagonism (I) between treatment and mechanical loading was assessed using the interaction term “treatment:loading” from a fitted linear model applied to Δw20-24.

| Readout | N | w15 | **w20 (baseline)** | w22 | w24 | **Δw20-24 [%]** | ○ | ○ | ○ | ○ | ○ | ● | ● | ● | Synergy |
|---|---|---|---|---|---|---|---|---|---|---|---|---|---|---|---|
| **VEH (0N: n=25; 8N: n=26)** | | | | | | | | | | | | | | | |
| BV/TV [%] | 0N | 18.900 ± 0.444 | 14.319 ± 0.416 | 12.801 ± 0.418 | 12.832 ± 0.372 | -10.051 ± 1.525 | x | *** <0.001 | *** <0.001 | *** <0.001 | *** <0.001 | *** <0.001 | *** <0.001 | *** <0.001 | - |
| | 8N | 19.650 ± 0.590 | 14.692 ± 0.306 | 15.761 ± 0.319 | 17.206 ± 0.343 | 17.652 ± 2.166 | *** <0.001 | x | 0.996 | 1.000 | 0.938 | 0.204 | *** <0.001 | *** <0.001 | - |
| Tb.Th [μm] | 0N | 74.916 ± 0.984 | 65.004 ± 0.691 | 63.889 ± 0.802 | 65.257 ± 0.724 | 0.471 ± 0.877 | x | *** <0.001 | 0.304 | ** 0.002 | *** <0.001 | *** <0.001 | *** <0.001 | *** <0.001 | - |
| | 8N | 76.215 ± 1.112 | 66.913 ± 0.735 | 73.023 ± 0.688 | 80.531 ± 0.909 | 20.606 ± 1.620 | *** <0.001 | x | *** <0.001 | *** <0.001 | 0.957 | 0.996 | *** <0.001 | *** <0.001 | - |
| Tb.N [1/mm] | 0N | 3.054 ± 0.066 | 2.990 ± 0.066 | 2.819 ± 0.070 | 2.727 ± 0.064 | -8.777 ± 0.803 | x | 0.985 | *** <0.001 | 0.384 | 0.823 | *** <0.001 | 0.487 | 1.000 | - |
| | 8N | 3.128 ± 0.065 | 2.975 ± 0.045 | 2.814 ± 0.049 | 2.736 ± 0.048 | -8.071 ± 0.663 | 0.985 | x | *** <0.001 | 0.817 | 0.395 | *** <0.001 | 0.880 | 0.998 | - |
| Conn.D [1/mm³] | 0N | 59.813 ± 2.222 | 62.528 ± 2.213 | 55.359 ± 2.327 | 48.863 ± 1.797 | -21.236 ± 2.242 | x | 0.187 | *** <0.001 | *** <0.001 | * 0.019 | 0.955 | 0.082 | *** <0.001 | - |
| | 8N | 62.461 ± 3.573 | 61.758 ± 1.815 | 51.572 ± 2.309 | 43.168 ± 1.661 | -29.454 ± 2.523 | 0.187 | x | *** <0.001 | *** <0.001 | 0.774 | 0.068 | 0.965 | *** <0.001 | - |
| Strength [N] | 0N | 35.872 ± 0.792 | 24.467 ± 0.830 | 23.863 ± 0.665 | 25.511 ± 0.597 | 6.010 ± 2.751 | x | *** <0.001 | *** <0.001 | *** <0.001 | *** <0.001 | *** <0.001 | *** <0.001 | *** <0.001 | - |
| | 8N | 35.551 ± 1.177 | 26.061 ± 0.723 | 30.414 ± 0.595 | 34.597 ± 0.538 | 34.657 ± 3.451 | *** <0.001 | x | 0.972 | 1.000 | 0.848 | 0.787 | *** <0.001 | *** <0.001 | - |
| SED [kPa] | 0N | 1.328 ± 0.045 | 2.369 ± 0.167 | 2.397 ± 0.138 | 2.152 ± 0.087 | -5.127 ± 3.305 | x | *** <0.001 | *** <0.001 | *** <0.001 | *** <0.001 | *** <0.001 | *** <0.001 | *** <0.001 | - |
| | 8N | 5.589 ± 0.347 | 8.298 ± 0.345 | 6.483 ± 0.184 | 5.287 ± 0.124 | -34.240 ± 2.278 | *** <0.001 | x | 0.991 | 0.999 | 0.908 | 0.156 | *** <0.001 | *** <0.001 | - |
| eff [με] | 0N | 386.583 ± 6.055 | 486.519 ± 13.443 | 490.780 ± 11.852 | 471.776 ± 8.528 | -2.334 ± 1.387 | x | *** <0.001 | *** <0.001 | *** <0.001 | *** <0.001 | *** <0.001 | *** <0.001 | *** <0.001 | - |
| | 8N | 777.540 ± 14.223 | 926.800 ± 17.599 | 836.932 ± 10.786 | 767.422 ± 8.612 | -16.700 ± 1.265 | *** <0.001 | x | 0.930 | 0.981 | 0.806 | * 0.047 | *** <0.001 | *** <0.001 | - |

| | | | | | | | | | | | | | | | |
|---|---|---|---|---|---|---|---|---|---|---|---|---|---|---|---|
| **BIS (0N: n=9; 8N: n=9)** | | | | | | | | | | | | | | | |
| BV/TV [%] | 0N | 17.249 ± 0.844 | 13.119 ± 0.381 | 13.965 ± 0.209 | 14.942 ± 0.224 | 14.448 ± 2.834 | *** <0.001 | 0.996 | x | 0.972 | 0.773 | 0.147 | *** <0.001 | *** <0.001 | - |
| | 8N | 18.046 ± 0.800 | 14.315 ± 0.538 | 15.827 ± 0.506 | 18.287 ± 0.564 | 28.040 ± 1.483 | *** <0.001 | 0.204 | 0.147 | 0.693 | 0.961 | x | *** <0.001 | *** <0.001 | **I* 0.029** |
| Tb.Th [µm] | 0N | 68.818 ± 1.542 | 63.445 ± 0.775 | 64.994 ± 0.933 | 66.507 ± 1.143 | 4.832 ± 1.334 | 0.304 | *** <0.001 | x | 0.873 | *** <0.001 | *** <0.001 | *** <0.001 | *** <0.001 | - |
| | 8N | 72.772 ± 1.571 | 65.685 ± 1.408 | 70.857 ± 1.423 | 78.318 ± 1.598 | 19.271 ± 0.832 | *** <0.001 | 0.996 | *** <0.001 | * 0.038 | 0.808 | x | *** <0.001 | *** <0.001 | 0.228 |
| Tb.N [1/mm] | 0N | 3.049 ± 0.091 | 2.900 ± 0.087 | 2.890 ± 0.082 | 2.929 ± 0.079 | 1.092 ± 0.968 | *** <0.001 | *** <0.001 | x | *** <0.001 | *** <0.001 | 0.563 | *** <0.001 | *** <0.001 | - |
| | 8N | 3.002 ± 0.075 | 2.928 ± 0.077 | 2.883 ± 0.086 | 2.887 ± 0.070 | -1.365 ± 0.722 | *** <0.001 | *** <0.001 | 0.563 | ** 0.001 | *** <0.001 | x | ** 0.001 | *** <0.001 | 0.059 |
| Conn.D [1/mm$^3$] | 0N | 73.848 ± 5.318 | 65.494 ± 3.108 | 66.346 ± 2.951 | 65.816 ± 3.452 | 1.111 ± 5.102 | *** <0.001 | *** <0.001 | x | 0.999 | *** <0.001 | * 0.037 | *** <0.001 | *** <0.001 | - |
| | 8N | 61.471 ± 4.498 | 66.525 ± 4.171 | 62.080 ± 4.280 | 54.486 ± 3.530 | -17.793 ± 2.654 | 0.955 | 0.068 | * 0.037 | 0.146 | ** 0.006 | x | * 0.025 | *** <0.001 | 0.115 |
| Strength [N] | 0N | 33.850 ± 1.639 | 24.078 ± 1.059 | 28.051 ± 0.608 | 30.169 ± 0.503 | 26.707 ± 4.422 | *** <0.001 | 0.972 | x | 0.957 | 0.485 | 0.421 | *** <0.001 | *** <0.001 | - |
| | 8N | 35.956 ± 1.330 | 25.134 ± 0.774 | 30.766 ± 0.844 | 35.722 ± 0.938 | 42.376 ± 1.905 | *** <0.001 | 0.787 | 0.421 | 0.966 | 1.000 | x | *** <0.001 | *** <0.001 | 0.195 |
| SED [kPa] | 0N | 1.444 ± 0.107 | 2.283 ± 0.154 | 1.778 ± 0.060 | 1.590 ± 0.055 | -28.601 ± 3.570 | *** <0.001 | 0.991 | x | 1.000 | 0.67 | 0.093 | *** <0.001 | *** <0.001 | - |
| | 8N | 5.121 ± 0.353 | 8.256 ± 0.364 | 5.893 ± 0.193 | 4.661 ± 0.147 | -43.254 ± 1.107 | *** <0.001 | 0.156 | 0.093 | 0.139 | 0.957 | x | 0.222 | *** <0.001 | **I* 0.031** |
| eff [µε] | 0N | 398.883 ± 14.998 | 478.677 ± 14.787 | 433.711 ± 7.577 | 413.977 ± 7.597 | -13.147 ± 1.754 | *** <0.001 | 0.930 | x | 1.000 | 0.342 | * 0.012 | *** <0.001 | *** <0.001 | - |
| | 8N | 752.617 ± 22.149 | 915.854 ± 15.201 | 790.326 ± 9.229 | 712.192 ± 8.389 | -22.168 ± 0.729 | *** <0.001 | * 0.047 | * 0.012 | * 0.020 | 0.903 | x | 0.054 | *** <0.001 | 0.121 |
| **PTH (0N: n=10; 8N: n=9)** | | | | | | | | | | | | | | | |
| BV/TV [%] | 0N | 23.586 ± 0.775 | 15.701 ± 0.685 | 16.515 ± 0.532 | 19.034 ± 0.585 | 22.126 ± 3.170 | *** <0.001 | 1.000 | 0.972 | x | 0.999 | 0.693 | *** <0.001 | *** <0.001 | - |
| | 8N | 24.396 ± 1.283 | 17.112 ± 0.514 | 20.879 ± 0.331 | 24.870 ± 0.672 | 46.320 ± 5.608 | *** <0.001 | *** <0.001 | *** <0.001 | *** <0.001 | *** <0.001 | *** <0.001 | x | 0.262 | 0.592 |
| Tb.Th [µm] | 0N | 81.480 ± 1.464 | 64.539 ± 1.290 | 65.570 ± 0.969 | 71.904 ± 1.165 | 11.579 ± 1.499 | ** 0.002 | *** <0.001 | 0.873 | x | *** <0.001 | * 0.038 | *** <0.001 | *** <0.001 | - |
| | 8N | 81.164 ± 3.793 | 65.286 ± 1.728 | 76.065 ± 1.834 | 86.769 ± 2.229 | 33.334 ± 3.763 | *** <0.001 | *** <0.001 | *** <0.001 | *** <0.001 | * 0.010 | *** <0.001 | x | *** <0.001 | 0.671 |
| Tb.N [1/mm] | 0N | 3.314 ± 0.067 | 3.182 ± 0.049 | 3.027 ± 0.051 | 2.981 ± 0.044 | -6.285 ± 0.925 | 0.384 | 0.817 | *** <0.001 | x | 0.059 | ** 0.001 | 1.000 | 0.673 | - |
| | 8N | 3.546 ± 0.092 | 3.386 ± 0.077 | 3.220 ± 0.084 | 3.172 ± 0.077 | -6.331 ± 0.842 | 0.487 | 0.880 | *** <0.001 | 1.000 | 0.086 | ** 0.001 | x | 0.739 | 0.605 |
| Conn.D [1/mm$^3$] | 0N | 53.684 ± 2.677 | 61.894 ± 2.880 | 63.813 ± 4.428 | 57.276 ± 4.012 | -6.361 ± 6.479 | *** <0.001 | *** <0.001 | 0.999 | x | *** <0.001 | 0.146 | *** <0.001 | *** <0.001 | - |
| | 8N | 63.366 ± 5.082 | 75.111 ± 3.933 | 56.492 ± 3.811 | 53.270 ± 3.905 | -28.998 ± 3.673 | 0.082 | 0.965 | *** <0.001 | *** <0.001 | 1.000 | * 0.025 | x | 0.100 | **I* 0.032** |
| Strength [N] | 0N | 40.605 ± 1.161 | 24.955 ± 1.130 | 28.662 ± 0.647 | 34.113 ± 0.598 | 38.630 ± 4.967 | *** <0.001 | 1.000 | 0.957 | x | 0.98 | 0.966 | *** <0.001 | *** <0.001 | - |
| | 8N | 42.442 ± 1.989 | 27.409 ± 1.160 | 37.233 ± 0.684 | 46.164 ± 0.789 | 70.600 ± 7.118 | *** <0.001 | *** <0.001 | *** <0.001 | *** <0.001 | *** <0.001 | *** <0.001 | x | * 0.021 | 0.692 |
| SED [kPa] | 0N | 1.136 ± 0.072 | 2.358 ± 0.185 | 1.938 ± 0.080 | 1.478 ± 0.056 | -35.309 ± 3.503 | *** <0.001 | 0.999 | 1.000 | x | 0.787 | 0.139 | *** <0.001 | *** <0.001 | - |
| | 8N | 4.250 ± 0.374 | 7.958 ± 0.515 | 4.920 ± 0.159 | 3.531 ± 0.112 | -54.392 ± 2.848 | *** <0.001 | *** <0.001 | *** <0.001 | *** <0.001 | * 0.011 | 0.222 | x | 0.557 | 0.126 |
| eff [µε] | 0N | 363.814 ± 10.526 | 497.278 ± 16.297 | 464.109 ± 9.004 | 411.071 ± 7.738 | -16.890 ± 1.919 | *** <0.001 | 0.981 | 1.000 | x | 0.463 | * 0.020 | *** <0.001 | *** <0.001 | - |
| | 8N | 699.888 ± 29.012 | 921.744 ± 25.290 | 752.274 ± 12.408 | 646.306 ± 11.010 | -29.526 ± 2.025 | *** <0.001 | *** <0.001 | *** <0.001 | *** <0.001 | *** <0.001 | 0.054 | x | 0.106 | 0.620 |
| **SclAB (0N: n=9; 8N: n=8)** | | | | | | | | | | | | | | | |
| BV/TV [%] | 0N | 23.456 ± 1.169 | 15.295 ± 0.805 | 16.158 ± 0.758 | 19.012 ± 0.853 | 25.329 ± 4.223 | *** <0.001 | 0.938 | 0.773 | 0.999 | x | 0.961 | *** <0.001 | *** <0.001 | - |
| | 8N | 23.176 ± 0.994 | 15.116 ± 0.518 | 18.935 ± 0.740 | 23.922 ± 0.834 | 59.408 ± 7.448 | *** <0.001 | *** <0.001 | *** <0.001 | *** <0.001 | *** <0.001 | *** <0.001 | 0.262 | x | 0.311 |

| | | | | | | | | | | | | | | | |
|---|---|---|---|---|---|---|---|---|---|---|---|---|---|---|---|
| Tb.Th [µm] | 0N | 79.495 ± 2.283 | 63.984 ± 1.509 | 69.422 ± 1.345 | 79.953 ± 1.374 | 25.268 ± 2.307 | *** <0.001 | 0.957 | *** <0.001 | *** <0.001 | x | 0.808 | * 0.010 | *** <0.001 | - |
| | 8N | 80.377 ± 2.471 | 62.352 ± 2.108 | 76.745 ± 2.556 | 93.350 ± 4.963 | 49.949 ± 6.862 | *** <0.001 | *** <0.001 | *** <0.001 | *** <0.001 | *** <0.001 | *** <0.001 | *** <0.001 | x | 0.297 |
| Tb.N [1/mm] | 0N | 3.328 ± 0.092 | 3.113 ± 0.090 | 2.906 ± 0.080 | 2.787 ± 0.074 | -10.424 ± 0.900 | 0.823 | 0.395 | *** <0.001 | 0.059 | x | *** <0.001 | 0.086 | 0.944 | - |
| | 8N | 3.366 ± 0.084 | 3.286 ± 0.112 | 3.081 ± 0.121 | 3.017 ± 0.122 | -8.310 ± 0.632 | 1.000 | 0.998 | *** <0.001 | 0.673 | 0.944 | *** <0.001 | 0.739 | x | 0.467 |
| Conn.D [1/mm$^3$] | 0N | 68.842 ± 4.411 | 68.475 ± 5.835 | 52.715 ± 3.706 | 43.315 ± 2.051 | -35.023 ± 2.766 | * 0.019 | 0.774 | *** <0.001 | *** <0.001 | x | ** 0.006 | 1.000 | 0.249 | - |
| | 8N | 58.353 ± 4.786 | 79.184 ± 11.397 | 53.348 ± 8.541 | 43.297 ± 6.892 | -45.407 ± 2.772 | *** <0.001 | *** <0.001 | *** <0.001 | *** <0.001 | 0.249 | *** <0.001 | 0.100 | x | 0.733 |
| Strength [N] | 0N | 39.602 ± 1.627 | 24.416 ± 0.906 | 28.509 ± 1.228 | 35.787 ± 1.268 | 47.028 ± 3.567 | *** <0.001 | 0.848 | 0.485 | 0.980 | x | 1.000 | *** <0.001 | *** <0.001 | - |
| | 8N | 41.085 ± 1.952 | 23.236 ± 1.413 | 33.566 ± 1.230 | 44.257 ± 1.330 | 95.209 ± 12.862 | *** <0.001 | *** <0.001 | *** <0.001 | *** <0.001 | *** <0.001 | *** <0.001 | * 0.021 | x | **S* 0.045** |
| SED [kPa] | 0N | 1.161 ± 0.066 | 2.378 ± 0.173 | 1.908 ± 0.124 | 1.318 ± 0.070 | -43.770 ± 2.342 | *** <0.001 | 0.908 | 0.670 | 0.787 | x | 0.957 | * 0.011 | *** <0.001 | - |
| | 8N | 4.492 ± 0.302 | 10.492 ± 1.156 | 5.623 ± 0.248 | 3.696 ± 0.187 | -62.794 ± 3.192 | *** <0.001 | *** <0.001 | *** <0.001 | *** <0.001 | *** <0.001 | *** <0.001 | 0.557 | x | 0.440 |
| eff [µε] | 0N | 366.189 ± 9.662 | 497.919 ± 17.193 | 454.488 ± 13.976 | 385.734 ± 10.069 | -22.275 ± 1.500 | *** <0.001 | 0.806 | 0.342 | 0.463 | x | 0.903 | *** <0.001 | *** <0.001 | - |
| | 8N | 722.750 ± 22.926 | 1030.171 ± 40.492 | 790.286 ± 17.042 | 654.367 ± 17.076 | -35.938 ± 2.566 | *** <0.001 | *** <0.001 | *** <0.001 | *** <0.001 | *** <0.001 | *** <0.001 | 0.106 | x | 0.859 |

**Table S.3:** Absolute values and %-changes in bone volume fraction (BV/TV) of the three vehicle groups of the four experiments (see Methods). The animals in Experiment 3 (SclAB) were supplied by a different provider; see Methods section "Study design". The VEH 0N groups received sham loading, the VEH 8N groups cyclic mechanical loading according to an established loading protocol[1] in the sixth caudal vertebra between week 20 and 24. Values are presented as group means ± standard error in weeks 15, 20, 22, and 24 (w15, w20, w22, w24), Δ represents the %-change from week 20 to 24 (Δw20-24). One-way ANOVA for the shown vehicle groups followed by Tukey's post-hoc test was used to calculate significant differences between groups. *p<0.05, **p<0.01, ***p<0.001.

| BV/TV [%] | w15 | **w20 (baseline)** | w22 | w24 | Δw20-24 [%] |
|---|---|---|---|---|---|
| **Analysis 1: VEH 0N (n = 25)** | | | | | |
| Experiment 1 | 17.398 ± 0.489 | 13.172 ± 0.741 | 11.467 ± 0.560 | 11.247 ± 0.524 | -13.840 ± 2.731 |
| Experiment 2 | 18.854 ± 0.650 | 14.204 ± 0.378 | 13.065 ± 0.310 | 13.387 ± 0.293 | -5.606 ± 1.390 |
| Experiment 3 | 20.636 ± 0.765 | 15.724 ± 0.716 | 14.036 ± 0.928 | 14.060 ± 0.615 | -10.232 ± 2.875 |
| P-value: 1 vs. 2; 2 vs. 3; 1 vs. 3 | 0.861; 0.673; ***0.014** | 0.987; 0.849; 0.130 | 0.773; 0.994; 0.124 | 0.350; 1.000; 0.060 | 0.064;0.408; 0.557 |
| **Analysis 2: VEH 8N (n = 26)** | | | | | |
| Experiment 1 | 16.998 ± 0.884 | 13.811 ± 0.733 | 14.369 ± 0.680 | 15.421 ± 0.635 | 12.388 ± 3.810 |
| Experiment 2 | 18.809 ± 0.699 | 14.714 ± 0.527 | 15.776 ± 0.431 | 17.231 ± 0.321 | 17.842 ± 3.044 |
| Experiment 3 | 22.179 ± 0.573 | 15.288 ± 0.324 | 16.721 ± 0.345 | 18.434 ± 0.405 | 21.166 ± 4.020 |
| P-value: 1 vs. 2; 2 vs. 3; 1 vs. 3 | 0.534; **0.001; *<0.001** | 0.992; 1.000; 0.7551 | 0.825; 0.976; 0.114 | 0.493; 0.878; ****0.010** | 0.588; 0.786; 0.252 |

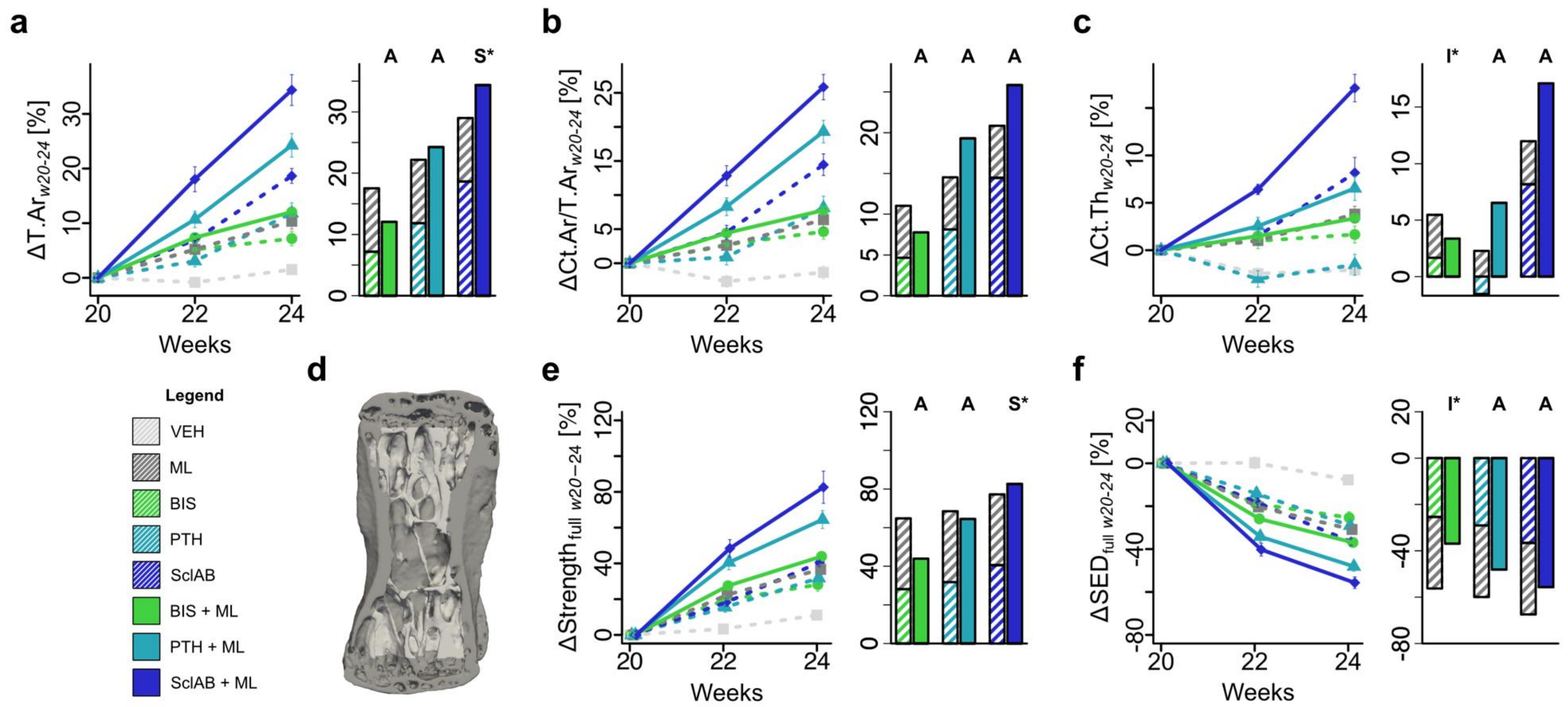


**Figure S.1: Mechanical loading increases the cortical effects of anabolic and anti-catabolic treatments, contributing to increased predicted strength. (a-f)** Changes from baseline. Data are presented as mean ± standard error. Each data point represents one animal (biological replicate; VEH: n=25, ML: n=26, BIS 0N: n=9, BIS 8N: n=9, PTH 0N: n=10, PTH 8N: n=9, SclAB 0N: n=9, SclAB 8N: n=8) i **(a)** total area (ΔT.Ar) **(b)** cortical area per total area (ΔCt.Ar/T.Ar) **(c)** cortical thickness (ΔCt.Th). **(d)** Visual representation of a representative cortical shell of a VEH 0N animal at week 20 **(e)** full predicted strength including both trabecular and cortical compartments (Δfull Strength), and **(f)** full strain energy density (Δfull SED). Bars show group means of sum of single vs. combined treatment indicating: antagonistic (I), additive (A) or synergistic (S) effects determined via the interaction term “treatment:loading” from a linear model on the %-change from week 20 to 24. Source data are provided as a Source Data file.

**Table S.4:** Cortical morphometric and full mechanical parameters of the non-loaded (0N) and loaded (8N) 6th caudal vertebrae. Values are given as group means ± standard error, Δ represents the %-change from week 20 to 24 (Δw20-24). ‘full’ denotes that both trabecular and cortical compartments were included. Two-way ANOVA over time for the shown animal groups followed by Tukey’s post-hoc test was used on Δw20-24. ‘x’ denotes the reference group; asterisks (*, **, ***) indicate groups that are significantly different from 'x'. Significance is indicated as: *p<0.05, **p<0.01, ***p<0.001. Synergy (S) or antagonism (I) between treatment and mechanical loading was assessed using the interaction term “treatment:loading” from a fitted linear model applied to Δw20-24.

| Readout | N | w15 | **w20 (baseline)** | w22 | w24 | **Δw20-24 [%]** | ○ | ○ | ○ | ○ | ○ | ● | ● | ● | Synergy |
|---|---|---|---|---|---|---|---|---|---|---|---|---|---|---|---|
| **VEH (0N: n=25; 8N: n=26)** | | | | | | | | | | | | | | | |
| Tt.Ar [mm$^2$] | 0N | 0.782 ± 0.011 | 0.690 ± 0.013 | 0.683 ± 0.013 | 0.699 ± 0.011 | 1.512 ± 0.906 | x | *** <0.001 | *** <0.001 | *** <0.001 | *** <0.001 | *** <0.001 | *** <0.001 | *** <0.001 | - |
| | 8N | 0.784 ± 0.016 | 0.719 ± 0.015 | 0.754 ± 0.012 | 0.789 ± 0.010 | 10.352 ± 1.435 | *** <0.001 | x | 0.941 | 1.000 | ** 0.008 | 0.861 | *** <0.001 | *** <0.001 | - |
| Ct.Ar/Tt.Ar [%] | 0N | 39.691 ± 0.437 | 33.846 ± 0.544 | 32.899 ± 0.507 | 33.319 ± 0.456 | -1.350 ± 0.923 | x | *** <0.001 | *** <0.001 | *** <0.001 | *** <0.001 | *** <0.001 | *** <0.001 | *** <0.001 | - |
| | 8N | 39.412 ± 0.669 | 34.560 ± 0.528 | 35.412 ± 0.444 | 36.651 ± 0.357 | 6.388 ± 1.185 | *** <0.001 | x | 0.997 | 1.000 | *** <0.001 | 0.876 | *** <0.001 | *** <0.001 | - |
| Ct.BS/BV [1/mm] | 0N | 10.299 ± 0.137 | 12.007 ± 0.211 | 12.140 ± 0.201 | 11.859 ± 0.159 | -0.991 ± 0.925 | x | *** <0.001 | 0.079 | *** <0.001 | *** <0.001 | *** <0.001 | *** <0.001 | *** <0.001 | - |
| | 8N | 10.402 ± 0.225 | 11.741 ± 0.200 | 11.319 ± 0.170 | 10.804 ± 0.134 | -7.654 ± 1.063 | *** <0.001 | x | 0.486 | *** <0.001 | * 0.011 | 1.000 | 1.000 | *** <0.001 | - |
| Ct.Th [µm] | 0N | 147.375 ± 1.513 | 138.136 ± 1.626 | 134.676 ± 1.629 | 135.171 ± 1.463 | -2.078 ± 0.571 | x | *** <0.001 | *** <0.001 | 1.000 | *** <0.001 | *** <0.001 | *** <0.001 | *** <0.001 | - |
| | 8N | 147.041 ± 2.279 | 140.147 ± 1.750 | 141.499 ± 1.610 | 145.254 ± 1.362 | 3.810 ± 0.829 | *** <0.001 | x | 0.892 | *** <0.001 | 0.076 | 1.000 | 0.181 | *** <0.001 | - |
| full Strength [N] | 0N | 40.461 ± 0.948 | 30.545 ± 0.962 | 31.360 ± 0.863 | 33.541 ± 0.715 | 11.041 ± 1.994 | x | *** <0.001 | *** <0.001 | *** <0.001 | *** <0.001 | *** <0.001 | *** <0.001 | *** <0.001 | - |
| | 8N | 39.320 ± 1.494 | 32.444 ± 0.871 | 39.392 ± 0.726 | 43.815 ± 0.594 | 36.657 ± 2.946 | *** <0.001 | x | 0.595 | 0.530 | 1.000 | 0.519 | *** <0.001 | *** <0.001 | - |
| full SED [kPa] | 0N | 0.926 ± 0.029 | 1.387 ± 0.087 | 1.368 ± 0.072 | 1.245 ± 0.049 | -7.748 ± 2.248 | x | *** <0.001 | *** <0.001 | *** <0.001 | *** <0.001 | *** <0.001 | *** <0.001 | *** <0.001 | - |
| | 8N | 4.085 ± 0.372 | 4.929 ± 0.202 | 3.860 ± 0.112 | 3.320 ± 0.074 | -30.863 ± 1.943 | *** <0.001 | x | 0.856 | 0.723 | 0.996 | 0.296 | *** <0.001 | *** <0.001 | - |
| full eff [µε] | 0N | 311.044 ± 4.443 | 365.964 ± 8.493 | 368.154 ± 8.102 | 355.648 ± 6.220 | -2.478 ± 0.914 | x | *** <0.001 | *** <0.001 | *** <0.001 | *** <0.001 | *** <0.001 | *** <0.001 | *** <0.001 | - |
| | 8N | 629.025 ± 14.398 | 699.091 ± 13.042 | 632.674 ± 8.494 | 592.658 ± 6.395 | -14.792 ± 1.083 | *** <0.001 | x | 0.695 | 0.479 | 0.996 | 0.273 | *** <0.001 | *** <0.001 | - |
| **BIS (0N: n=9; 8N: n=9)** | | | | | | | | | | | | | | | |
| Tt.Ar [mm$^2$] | 0N | 0.765 ± 0.023 | 0.691 ± 0.023 | 0.725 ± 0.018 | 0.738 ± 0.018 | 7.178 ± 1.296 | *** <0.001 | 0.941 | x | 0.994 | ** 0.003 | 0.420 | *** <0.001 | *** <0.001 | - |
| | 8N | 0.818 ± 0.020 | 0.727 ± 0.012 | 0.781 ± 0.011 | 0.815 ± 0.011 | 12.051 ± 0.579 | *** <0.001 | 0.861 | 0.420 | 0.866 | 0.603 | x | *** <0.001 | *** <0.001 | 0.221 |
| Ct.Ar/Tt.Ar [%] | 0N | 37.553 ± 0.705 | 33.151 ± 0.803 | 34.010 ± 0.609 | 34.627 ± 0.562 | 4.640 ± 1.113 | *** <0.001 | 0.997 | x | 0.999 | ** 0.002 | 0.697 | *** <0.001 | *** <0.001 | - |
| | 8N | 38.966 ± 0.890 | 33.675 ± 0.616 | 35.157 ± 0.579 | 36.279 ± 0.592 | 7.769 ± 0.481 | *** <0.001 | 0.876 | 0.697 | 0.949 | 0.262 | x | *** <0.001 | *** <0.001 | 0.094 |
| Ct.BS/BV [1/mm] | 0N | 10.714 ± 0.230 | 12.065 ± 0.343 | 11.748 ± 0.255 | 11.553 ± 0.218 | -4.017 ± 1.282 | 0.079 | 0.486 | x | *** <0.001 | *** <0.001 | 0.925 | 0.561 | *** <0.001 | - |
| | 8N | 10.232 ± 0.286 | 11.630 ± 0.232 | 11.257 ± 0.250 | 10.854 ± 0.240 | -6.697 ± 0.451 | *** <0.001 | 1.000 | 0.925 | *** <0.001 | * 0.022 | x | 0.998 | *** <0.001 | 0.133 |
| Ct.Th [µm] | 0N | 140.764 ± 2.272 | 135.074 ± 2.399 | 136.251 ± 1.431 | 137.167 ± 1.609 | 1.655 ± 0.880 | *** <0.001 | 0.892 | x | ** 0.008 | * 0.014 | 0.958 | * 0.036 | *** <0.001 | - |

| | | | | | | | | | | | | | | | |
|---|---|---|---|---|---|---|---|---|---|---|---|---|---|---|---|
| | 8N | 147.476 ± 2.926 | 139.921 ± 1.724 | 142.026 ± 1.953 | 144.650 ± 2.047 | 3.363 ± 0.299 | *** <0.001 | 1.000 | 0.958 | *** <0.001 | 0.261 | x | 0.433 | *** <0.001 | I* 0.037 |
| Full Strength [N] | 0N | 39.812 ± 1.909 | 30.644 ± 1.358 | 36.288 ± 0.857 | 38.908 ± 0.800 | 28.138 ± 3.668 | *** <0.001 | 0.595 | x | 1.000 | 0.824 | * 0.043 | *** <0.001 | *** <0.001 | - |
| | 8N | 43.277 ± 1.518 | 32.238 ± 0.850 | 41.043 ± 0.898 | 46.341 ± 1.114 | 43.917 ± 1.870 | *** <0.001 | 0.519 | * 0.043 | * 0.032 | 0.726 | x | *** <0.001 | *** <0.001 | 0.195 |
| Full SED [kPa] | 0N | 0.984 ± 0.066 | 1.370 ± 0.097 | 1.095 ± 0.047 | 1.002 ± 0.038 | -25.369 ± 2.774 | *** <0.001 | 0.856 | x | 1.000 | 0.650 | 0.052 | *** <0.001 | *** <0.001 | - |
| | 8N | 3.369 ± 0.187 | 4.774 ± 0.204 | 3.528 ± 0.125 | 3.005 ± 0.114 | -36.924 ± 0.864 | *** <0.001 | 0.296 | 0.052 | * 0.026 | 0.905 | x | * 0.036 | *** <0.001 | I* 0.023 |
| Full eff [µε] | 0N | 321.049 ± 10.620 | 369.008 ± 12.270 | 337.400 ± 7.380 | 324.857 ± 6.547 | -11.628 ± 1.380 | *** <0.001 | 0.695 | x | 1.000 | 0.487 | * 0.023 | *** <0.001 | *** <0.001 | - |
| | 8N | 597.436 ± 14.782 | 689.779 ± 12.362 | 608.455 ± 9.457 | 566.119 ± 9.505 | -17.906 ± 0.438 | *** <0.001 | 0.273 | * 0.023 | ** 0.009 | 0.893 | x | ** 0.002 | *** <0.001 | I* 0.020 |
| **PTH (0N: n=10; 8N: n=9)** | | | | | | | | | | | | | | | |
| Tt.Ar [mm$^2$] | 0N | 0.804 ± 0.013 | 0.662 ± 0.013 | 0.682 ± 0.011 | 0.739 ± 0.013 | 11.826 ± 1.881 | *** <0.001 | 1.000 | 0.994 | x | * 0.028 | 0.866 | *** <0.001 | *** <0.001 | - |
| | 8N | 0.833 ± 0.033 | 0.714 ± 0.022 | 0.789 ± 0.018 | 0.885 ± 0.021 | 24.255 ± 2.152 | *** <0.001 | *** <0.001 | *** <0.001 | *** <0.001 | 0.103 | *** <0.001 | x | *** <0.001 | 0.242 |
| Ct.Ar/Tt.Ar [%] | 0N | 42.335 ± 0.929 | 33.494 ± 0.908 | 33.730 ± 0.705 | 36.115 ± 0.691 | 8.132 ± 1.694 | *** <0.001 | 1.000 | 0.999 | x | * 0.01 | 0.949 | *** <0.001 | *** <0.001 | - |
| | 8N | 42.568 ± 1.192 | 34.894 ± 0.967 | 37.733 ± 0.853 | 41.565 ± 0.973 | 19.310 ± 1.651 | *** <0.001 | *** <0.001 | *** <0.001 | *** <0.001 | 0.058 | *** <0.001 | x | ** 0.005 | 0.198 |
| Ct.BS/BV [1/mm] | 0N | 9.778 ± 0.207 | 12.278 ± 0.267 | 13.043 ± 0.248 | 12.840 ± 0.272 | 4.720 ± 1.754 | *** <0.001 | *** <0.001 | *** <0.001 | x | *** <0.001 | *** <0.001 | *** <0.001 | *** <0.001 | - |
| | 8N | 9.600 ± 0.419 | 12.006 ± 0.230 | 11.595 ± 0.169 | 10.972 ± 0.164 | -8.509 ± 1.020 | *** <0.001 | 1.000 | 0.561 | *** <0.001 | 0.140 | 0.998 | x | *** <0.001 | I* 0.010 |
| Ct.Th [µm] | 0N | 154.496 ± 2.629 | 136.411 ± 2.520 | 132.155 ± 1.894 | 134.166 ± 2.007 | -1.537 ± 1.109 | 1.000 | *** <0.001 | ** 0.008 | x | *** <0.001 | *** <0.001 | *** <0.001 | *** <0.001 | - |
| | 8N | 157.820 ± 5.308 | 141.347 ± 3.669 | 144.761 ± 2.963 | 150.293 ± 2.699 | 6.526 ± 1.264 | *** <0.001 | 0.181 | * 0.036 | *** <0.001 | 1.000 | 0.433 | x | *** <0.001 | 0.259 |
| Full Strength [N] | 0N | 42.623 ± 0.877 | 30.240 ± 1.049 | 34.705 ± 0.752 | 39.581 ± 0.838 | 31.813 ± 3.609 | *** <0.001 | 0.530 | 1.000 | x | 0.795 | * 0.032 | *** <0.001 | *** <0.001 | - |
| | 8N | 44.699 ± 1.587 | 32.872 ± 1.042 | 45.930 ± 0.831 | 53.723 ± 0.879 | 64.505 ± 4.979 | *** <0.001 | *** <0.001 | *** <0.001 | *** <0.001 | *** <0.001 | *** <0.001 | x | * 0.027 | 0.297 |
| Full SED [kPa] | 0N | 0.832 ± 0.030 | 1.374 ± 0.083 | 1.164 ± 0.044 | 0.959 ± 0.036 | -29.069 ± 2.878 | *** <0.001 | 0.723 | 1.000 | x | 0.516 | * 0.026 | *** <0.001 | *** <0.001 | - |
| | 8N | 3.152 ± 0.247 | 4.709 ± 0.265 | 3.058 ± 0.110 | 2.406 ± 0.082 | -48.096 ± 2.249 | *** <0.001 | *** <0.001 | *** <0.001 | *** <0.001 | *** <0.001 | * 0.036 | x | 0.385 | 0.420 |
| Full eff [µε] | 0N | 295.311 ± 5.349 | 367.309 ± 9.223 | 345.724 ± 6.331 | 316.393 ± 5.798 | -13.639 ± 1.487 | *** <0.001 | 0.479 | 1.000 | x | 0.324 | ** 0.009 | *** <0.001 | *** <0.001 | - |
| | 8N | 572.588 ± 21.462 | 684.048 ± 18.123 | 570.847 ± 10.029 | 510.037 ± 8.700 | -25.182 ± 1.561 | *** <0.001 | *** <0.001 | *** <0.001 | *** <0.001 | *** <0.001 | ** 0.002 | x | 0.112 | 0.781 |
| **SclAB (0N: n=9; 8N: n=8)** | | | | | | | | | | | | | | | |
| Tt.Ar [mm$^2$] | 0N | 0.782 ± 0.023 | 0.658 ± 0.015 | 0.703 ± 0.014 | 0.779 ± 0.011 | 18.645 ± 1.355 | *** <0.001 | ** 0.008 | ** 0.003 | * 0.028 | x | 0.603 | 0.103 | *** <0.001 | - |
| | 8N | 0.824 ± 0.027 | 0.658 ± 0.024 | 0.774 ± 0.022 | 0.881 ± 0.026 | 34.375 ± 2.824 | *** <0.001 | *** <0.001 | *** <0.001 | *** <0.001 | *** <0.001 | *** <0.001 | *** <0.001 | x | S* 0.034 |
| Ct.Ar/Tt.Ar [%] | 0N | 42.157 ± 1.502 | 34.044 ± 0.972 | 35.565 ± 0.975 | 38.893 ± 0.900 | 14.461 ± 1.580 | *** <0.001 | *** <0.001 | ** 0.002 | * 0.01 | x | 0.262 | 0.058 | *** <0.001 | - |
| | 8N | 43.400 ± 1.082 | 33.324 ± 1.467 | 37.466 ± 1.290 | 41.798 ± 1.506 | 25.845 ± 1.852 | *** <0.001 | *** <0.001 | *** <0.001 | *** <0.001 | *** <0.001 | *** <0.001 | ** 0.005 | x | 0.191 |
| Ct.BS/BV [1/mm] | 0N | 9.836 ± 0.394 | 12.183 ± 0.351 | 11.627 ± 0.282 | 10.322 ± 0.194 | -14.931 ± 1.937 | *** <0.001 | * 0.011 | *** <0.001 | *** <0.001 | x | * 0.022 | 0.140 | *** <0.001 | - |
| | 8N | 9.677 ± 0.315 | 12.657 ± 0.632 | 11.352 ± 0.646 | 9.819 ± 0.679 | -22.764 ± 1.593 | *** <0.001 | *** <0.001 | *** <0.001 | *** <0.001 | *** <0.001 | *** <0.001 | *** <0.001 | x | 0.641 |
| Ct.Th [µm] | 0N | 155.554 ± 5.396 | 137.670 ± 3.588 | 139.594 ± 3.126 | 148.587 ± 2.708 | 8.170 ± 1.617 | *** <0.001 | 0.076 | * 0.014 | *** <0.001 | x | 0.261 | 1.000 | *** <0.001 | - |

| | | | | | | | | | | | | | | | |
|---|---|---|---|---|---|---|---|---|---|---|---|---|---|---|---|
| | 8N | 158.253 ± 4.356 | 136.541 ± 4.262 | 145.251 ± 4.391 | 159.885 ± 5.154 | 17.105 ± 1.448 | *** <0.001 | *** <0.001 | *** <0.001 | *** <0.001 | *** <0.001 | *** <0.001 | *** <0.001 | x | 0.131 |
| Full Strength [N] | 0N | 40.739 ± 1.673 | 29.305 ± 0.663 | 34.671 ± 1.123 | 41.160 ± 1.033 | 40.547 ± 2.331 | *** <0.001 | 1.000 | 0.824 | 0.795 | x | 0.726 | *** <0.001 | *** <0.001 | - |
| | 8N | 45.012 ± 1.838 | 29.215 ± 1.573 | 42.882 ± 1.414 | 52.445 ± 1.272 | 82.593 ± 8.998 | *** <0.001 | *** <0.001 | *** <0.001 | *** <0.001 | *** <0.001 | *** <0.001 | * 0.027 | x | **S* 0.026** |
| Full SED [kPa] | 0N | 0.908 ± 0.059 | 1.424 ± 0.055 | 1.164 ± 0.057 | 0.899 ± 0.030 | -36.580 ± 1.702 | *** <0.001 | 0.996 | 0.650 | 0.516 | x | 0.905 | *** <0.001 | *** <0.001 | - |
| | 8N | 3.159 ± 0.190 | 5.730 ± 0.466 | 3.341 ± 0.134 | 2.467 ± 0.099 | -55.681 ± 2.691 | *** <0.001 | *** <0.001 | *** <0.001 | *** <0.001 | *** <0.001 | *** <0.001 | 0.385 | x | 0.448 |
| Full eff [με] | 0N | 306.378 ± 8.987 | 374.404 ± 6.860 | 344.760 ± 7.367 | 307.143 ± 4.588 | -17.873 ± 1.068 | *** <0.001 | 0.996 | 0.487 | 0.324 | x | 0.893 | *** <0.001 | *** <0.001 | - |
| | 8N | 577.451 ± 16.650 | 738.907 ± 20.446 | 591.704 ± 11.876 | 514.655 ± 11.210 | -30.134 ± 1.686 | *** <0.001 | *** <0.001 | *** <0.001 | *** <0.001 | *** <0.001 | *** <0.001 | 0.112 | x | 0.990 |

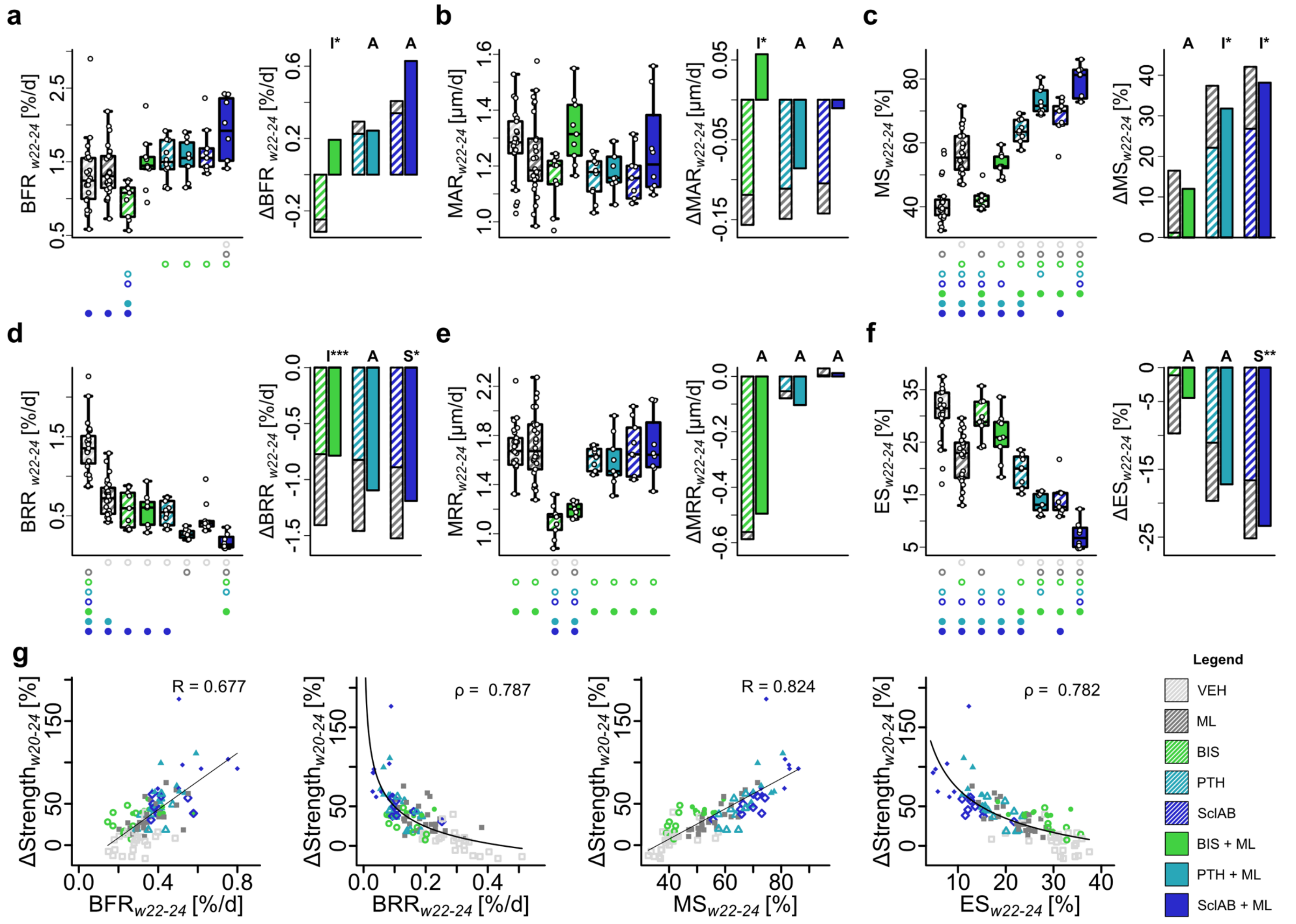
a
b
c
d
e
f
g
BFR w22-24 [%/d]
ΔBFR w22-24 [%/d]
MAR w22-24 [µm/d]
ΔMAR w22-24 [µm/d]
MS w22-24 [%]
ΔMS w22-24 [%]
BRR w22-24 [%/d]
ΔBRR w22-24 [%/d]
MRR w22-24 [µm/d]
ΔMRR w22-24 [µm/d]
ES w22-24 [%]
ΔES w22-24 [%]
I*
A
I***
S*
S**
ΔStrength w20-24 [%]
R = 0.677
ρ = 0.787
R = 0.824
ρ = 0.782
BFR w22-24 [%/d]
BRR w22-24 [%/d]
MS w22-24 [%]
ES w22-24 [%]
Legend
VEH
ML
BIS
PTH
SclAB
BIS + ML
PTH + ML
SclAB + ML

**Figure S.2: Time interval w22-24: Mechanical loading increases bone formation and decreases resorption rates of anabolic and anti-catabolic pharmacological treatments which contribute to increased predicted strength. (a)** bone formation rate per bone volume (BFR) **(b)** mineral apposition rate (MAR) **(c)** mineralizing surface per bone surface (MS) **(d)** bone resorption rate per bone volume (BRR) **(e)** mineral resorption rate (MRR) **(f)** eroded surface per bone surface (ES). **(a-f)** Each data point represents one animal (biological replicate; VEH: n=25, ML: n=26, BIS 0N: n=9, BIS 8N: n=9, PTH 0N: n=10, PTH 8N: n=9, SclAB 0N: n=9, SclAB 8N: n=8). The box spans the interquartile range (IQR) from the 25th to the 75th percentile. The median is shown as a line inside the box. Whiskers extend to the most extreme data points within 1.5 × IQR, points beyond this range are plotted as outliers. Δ denotes difference from VEH as no baseline value for dynamic bone morphometry rates was available. Bars show group means of sum of single vs. combined treatment indicating: antagonistic (I), additive (A) or synergistic (S) effects determined via the interaction term “treatment:loading” in a linear model on the $\log_2$-transformed values. Significances were calculated using one-way ANOVA followed by Tukey’s post-hoc test; significant differences are depicted with a symbol in the color of the corresponding group (o: to monotreatment, ●: to combined treatment, see Supplementary Table S.1 for color encoding, p<0.05). **(g)** Linear regression of ΔStrength with MS and BFR, power-law regression with ES and BRR in week interval w22-24. Source data are provided as a Source Data file.

**Table S.5:** Dynamic morphometric parameters of the non-loaded (0N) and loaded (8N) trabecular compartments of 6th caudal vertebrae, given as group means ± standard error in week intervals w20-22 and w22-24. The %-change between week intervals w20-22 and w22-24, calculated per animal, is depicted as well. One-way ANOVA in each time interval for the shown animal groups followed by Tukey’s post-hoc test was used on the absolute values. ‘w20-22’ or ‘w22-24’ denotes the reference group; asterisks (*, **, ***) indicate groups that are significantly different from it. Significance is indicated as: *p<0.05, **p<0.01, ***p<0.001. Synergy (S) or antagonism (I) between treatment and mechanical loading was assessed using the interaction term “treatment:loading” from a fitted linear model applied to the $\log_2$-transformed absolute values within each interval.

| Readout | N | w20-22 | w22-24 | %-change between w20-22 and w22-24 [%] | o | o | o | o | o | • | • | • | Synergy w20-22 | Synergy w22-24 |
|---|---|---|---|---|---|---|---|---|---|---|---|---|---|---|
| **VEH (0N: n=25; 8N: n=26)** | | | | | | | | | | | | | | |
| BFR [%/d] | 0N | 1.081 ± 0.064 | 1.300 ± 0.092 | 29.102 ± 13.091 | w20-22 | ** 0.008 | 0.767 | * 0.035 | 0.467 | * 0.022 | *** <0.001 | *** <0.001 | - | - |
| | | | | | w22-24 | 0.998 | 0.345 | 0.715 | 0.256 | 0.872 | 0.675 | ** 0.001 | | |
| | 8N | 1.580 ± 0.079 | 1.367 ± 0.067 | -9.603 ± 5.814 | ** 0.008 | w20-22 | 0.956 | 1.000 | 0.998 | 0.996 | ** 0.001 | *** <0.001 | - | - |
| | | | | | 0.998 | w22-24 | 0.130 | 0.939 | 0.536 | 0.986 | 0.915 | ** 0.006 | | |
| BRR [%/d] | 0N | 2.044 ± 0.116 | 1.365 ± 0.065 | -30.613 ± 3.390 | w20-22 | *** <0.001 | *** <0.001 | ** 0.01 | *** <0.001 | *** <0.001 | *** <0.001 | *** <0.001 | - | - |
| | | | | | w22-24 | *** <0.001 | *** <0.001 | *** <0.001 | *** <0.001 | *** <0.001 | *** <0.001 | *** <0.001 | | |
| | 8N | 1.111 ± 0.056 | 0.730 ± 0.045 | -32.952 ± 3.300 | *** <0.001 | w20-22 | 0.945 | 0.052 | 0.997 | 0.189 | * 0.019 | ** 0.007 | - | - |
| | | | | | *** <0.001 | w22-24 | 0.790 | 0.374 | 0.100 | 0.700 | *** <0.001 | *** <0.001 | | |
| MAR [µm/d] | 0N | 1.307 ± 0.038 | 1.270 ± 0.026 | -1.866 ± 2.070 | w20-22 | 0.702 | 1.000 | 0.859 | 0.952 | 0.120 | * 0.033 | * 0.046 | - | - |
| | | | | | w22-24 | 0.962 | 0.242 | 0.276 | 0.403 | 0.940 | 0.657 | 1.000 | | |

| | | | | | | | | | | | | | | |
|---|---|---|---|---|---|---|---|---|---|---|---|---|---|---|
| | 8N | 1.378 ± 0.025 | 1.233 ± 0.029 | -10.313 ± 1.841 | 0.702 | w20-22 | 0.984 | 1.000 | 0.285 | 0.778 | 0.457 | 0.500 | - | - |
| | | | | | 0.962 | w22-24 | 0.711 | 0.770 | 0.869 | 0.525 | 0.976 | 0.999 | | |
| MRR [µm/d] | 0N | 2.090 ± 0.048 | 1.691 ± 0.039 | -18.593 ± 1.820 | w20-22 | ** 0.01 | *** <0.001 | 0.990 | 0.664 | *** <0.001 | *** <0.001 | ** 0.001 | - | - |
| | | | | | w22-24 | 1.000 | *** <0.001 | 0.971 | 1.000 | *** <0.001 | 0.902 | 1.000 | | |
| | 8N | 1.905 ± 0.035 | 1.716 ± 0.052 | -9.932 ± 2.175 | ** 0.01 | w20-22 | *** <0.001 | 0.554 | 0.986 | *** <0.001 | 0.146 | 0.656 | - | - |
| | | | | | 1.000 | w22-24 | *** <0.001 | 0.877 | 1.000 | *** <0.001 | 0.743 | 1.000 | | |
| MS [%] | 0N | 35.367 ± 0.953 | 41.220 ± 1.269 | 19.303 ± 6.190 | w20-22 | *** <0.001 | * 0.02 | *** <0.001 | *** <0.001 | *** <0.001 | *** <0.001 | *** <0.001 | - | - |
| | | | | | w22-24 | *** <0.001 | 0.999 | *** <0.001 | *** <0.001 | *** <0.001 | *** <0.001 | *** <0.001 | | |
| | 8N | 50.426 ± 1.472 | 56.504 ± 1.328 | 12.979 ± 2.204 | *** <0.001 | w20-22 | * 0.042 | 0.991 | 1.000 | 1.000 | *** <0.001 | *** <0.001 | - | - |
| | | | | | *** <0.001 | w22-24 | *** <0.001 | * 0.028 | *** <0.001 | 0.792 | *** <0.001 | *** <0.001 | | |
| ES [%] | 0N | 33.408 ± 0.653 | 30.602 ± 0.974 | -7.816 ± 3.031 | w20-22 | *** <0.001 | 0.958 | 0.569 | *** <0.001 | * 0.045 | *** <0.001 | *** <0.001 | - | - |
| | | | | | w22-24 | *** <0.001 | 0.996 | *** <0.001 | *** <0.001 | 0.105 | *** <0.001 | *** <0.001 | | |
| | 8N | 24.911 ± 1.149 | 22.035 ± 0.894 | -10.109 ± 2.446 | *** <0.001 | w20-22 | ** 0.009 | 0.054 | 1.000 | 0.736 | ** 0.005 | *** <0.001 | - | - |
| | | | | | *** <0.001 | w22-24 | *** <0.001 | 0.717 | *** <0.001 | 0.177 | *** <0.001 | *** <0.001 | | |
| **BIS (0N: n=9; 8N: n=9)** | | | | | | | | | | | | | | |
| BFR [%/d] | 0N | 1.375 ± 0.063 | 0.984 ± 0.083 | -27.677 ± 6.537 | 0.767 | 0.956 | w20-22 | 0.897 | 1.000 | 0.807 | *** <0.001 | *** <0.001 | - | - |
| | | | | | 0.345 | 0.130 | w22-24 | * 0.034 | ** 0.006 | 0.072 | * 0.032 | *** <0.001 | | |
| | 8N | 1.716 ± 0.156 | 1.493 ± 0.124 | -10.502 ± 6.308 | * 0.022 | 0.996 | 0.807 | 1.000 | 0.952 | w20-22 | 0.085 | * 0.033 | 0.183 | **I* 0.018** |
| | | | | | 0.872 | 0.986 | 0.072 | 1.000 | 0.990 | w22-24 | 1.000 | 0.227 | | |
| BRR [%/d] | 0N | 0.947 ± 0.120 | 0.592 ± 0.079 | -36.308 ± 5.376 | *** <0.001 | 0.945 | w20-22 | * 0.018 | 0.800 | 0.943 | 0.564 | 0.346 | - | - |
| | | | | | *** <0.001 | 0.790 | w22-24 | 1.000 | 0.962 | 1.000 | 0.077 | ** 0.009 | | |
| | 8N | 0.745 ± 0.108 | 0.578 ± 0.068 | -18.239 ± 7.643 | *** <0.001 | 0.189 | 0.943 | *** <0.001 | 0.147 | w20-22 | 0.996 | 0.954 | **I* 0.017** | **I*** <0.001** |
| | | | | | *** <0.001 | 0.700 | 1.000 | 1.000 | 0.981 | w22-24 | 0.104 | * 0.013 | | |
| MAR [µm/d] | 0N | 1.325 ± 0.025 | 1.151 ± 0.033 | -12.895 ± 2.740 | 1.000 | 0.984 | w20-22 | 0.987 | 0.940 | 0.472 | 0.239 | 0.264 | - | - |
| | | | | | 0.242 | 0.711 | w22-24 | 1.000 | 1.000 | 0.072 | 0.999 | 0.643 | | |
| | 8N | 1.469 ± 0.048 | 1.328 ± 0.041 | -9.312 ± 2.371 | 0.120 | 0.778 | 0.472 | 0.932 | * 0.039 | w20-22 | 1.000 | 1.000 | 0.531 | **I** 0.002** |
| | | | | | 0.940 | 0.525 | 0.072 | 0.083 | 0.130 | w22-24 | 0.253 | 0.955 | | |
| MRR [µm/d] | 0N | 1.262 ± 0.045 | 1.104 ± 0.045 | -12.340 ± 2.781 | *** <0.001 | *** <0.001 | w20-22 | *** <0.001 | *** <0.001 | 1.000 | *** <0.001 | *** <0.001 | - | - |
| | | | | | *** <0.001 | *** <0.001 | w22-24 | *** <0.001 | *** <0.001 | 0.981 | *** <0.001 | *** <0.001 | | |
| | 8N | 1.234 ± 0.041 | 1.195 ± 0.019 | -2.511 ± 2.853 | *** <0.001 | *** <0.001 | 1.000 | *** <0.001 | *** <0.001 | w20-22 | *** <0.001 | *** <0.001 | 0.191 | 0.176 |
| | | | | | *** <0.001 | *** <0.001 | 0.981 | *** <0.001 | *** <0.001 | w22-24 | ** 0.003 | *** <0.001 | | |
| MS [%] | 0N | 43.210 ± 1.460 | 42.389 ± 1.092 | -1.195 ± 3.476 | * 0.02 | * 0.042 | w20-22 | 0.494 | 0.128 | 0.373 | *** <0.001 | *** <0.001 | - | - |

| | | | | | | | | | | | | | | |
|---|---|---|---|---|---|---|---|---|---|---|---|---|---|---|
| | | | | | 0.999 | *** <0.001 | w22-24 | *** <0.001 | *** <0.001 | ** 0.002 | *** <0.001 | *** <0.001 | | |
| | 8N | 49.278 ± 1.266 | 53.222 ± 1.160 | 8.439 ± 3.108 | *** <0.001 | 1.000 | 0.373 | 1.000 | 0.999 | w20-22 | *** <0.001 | *** <0.001 | I** 0.002 | 0.121 |
| | | | | | *** <0.001 | 0.792 | ** 0.002 | ** 0.004 | *** <0.001 | w22-24 | *** <0.001 | *** <0.001 | | |
| ES [%] | 0N | 31.464 ± 1.830 | 29.459 ± 1.287 | -5.278 ± 3.625 | 0.958 | ** 0.009 | w20-22 | 0.999 | * 0.024 | 0.694 | *** <0.001 | *** <0.001 | - | - |
| | | | | | 0.996 | *** <0.001 | w22-24 | *** <0.001 | *** <0.001 | 0.671 | *** <0.001 | *** <0.001 | | |
| | 8N | 27.798 ± 1.385 | 26.123 ± 1.562 | -5.820 ± 4.220 | * 0.045 | 0.736 | 0.694 | 0.950 | 0.711 | w20-22 | *** <0.001 | *** <0.001 | 0.075 | 0.067 |
| | | | | | 0.105 | 0.177 | 0.671 | * 0.015 | *** <0.001 | w22-24 | *** <0.001 | *** <0.001 | | |
| **PTH (0N: n=10; 8N: n=9)** | | | | | | | | | | | | | | |
| BFR [%/d] | 0N | 1.664 ± 0.140 | 1.526 ± 0.084 | -4.375 ± 7.003 | * 0.035 | 1.000 | 0.897 | w20-22 | 0.984 | 1.000 | * 0.037 | * 0.013 | - | - |
| | | | | | 0.715 | 0.939 | * 0.034 | w22-24 | 0.997 | 1.000 | 1.000 | 0.291 | | |
| | 8N | 2.374 ± 0.235 | 1.543 ± 0.088 | -30.211 ± 7.468 | *** <0.001 | ** 0.001 | *** <0.001 | * 0.037 | ** 0.003 | 0.085 | w20-22 | 1.000 | 0.741 | 0.635 |
| | | | | | 0.675 | 0.915 | * 0.032 | 1.000 | 0.999 | 1.000 | w22-24 | 0.379 | | |
| BRR [%/d] | 0N | 1.537 ± 0.078 | 0.541 ± 0.048 | -64.449 ± 3.282 | ** 0.01 | 0.052 | * 0.018 | w20-22 | 0.547 | *** <0.001 | *** <0.001 | *** <0.001 | - | - |
| | | | | | *** <0.001 | 0.374 | 1.000 | w22-24 | 0.999 | 1.000 | 0.194 | * 0.028 | | |
| | 8N | 0.617 ± 0.049 | 0.268 ± 0.020 | -55.085 ± 4.167 | *** <0.001 | * 0.019 | 0.564 | *** <0.001 | * 0.021 | 0.996 | w20-22 | 1.000 | S* 0.026 | 0.814 |
| | | | | | *** <0.001 | *** <0.001 | 0.077 | 0.194 | 0.576 | 0.104 | w22-24 | 0.991 | | |
| MAR [µm/d] | 0N | 1.386 ± 0.035 | 1.159 ± 0.024 | -16.012 ± 2.310 | 0.859 | 1.000 | 0.987 | w20-22 | 0.437 | 0.932 | 0.743 | 0.761 | - | - |
| | | | | | 0.276 | 0.770 | 1.000 | w22-24 | 1.000 | 0.083 | 1.000 | 0.698 | | |
| | 8N | 1.498 ± 0.041 | 1.185 ± 0.025 | -20.591 ± 2.008 | * 0.033 | 0.457 | 0.239 | 0.743 | * 0.012 | 1.000 | w20-22 | 1.000 | 0.836 | 0.361 |
| | | | | | 0.657 | 0.976 | 0.999 | 1.000 | 1.000 | 0.253 | w22-24 | 0.922 | | |
| MRR [µm/d] | 0N | 2.033 ± 0.037 | 1.612 ± 0.029 | -20.505 ± 1.824 | 0.990 | 0.554 | *** <0.001 | w20-22 | 0.993 | *** <0.001 | ** 0.006 | 0.068 | - | - |
| | | | | | 0.971 | 0.877 | *** <0.001 | w22-24 | 0.989 | *** <0.001 | 1.000 | 0.984 | | |
| | 8N | 1.719 ± 0.073 | 1.587 ± 0.069 | -6.995 ± 3.963 | *** <0.001 | 0.146 | *** <0.001 | ** 0.006 | 0.08 | *** <0.001 | w20-22 | 0.998 | 0.128 | 0.727 |
| | | | | | 0.902 | 0.743 | *** <0.001 | 1.000 | 0.957 | ** 0.003 | w22-24 | 0.945 | | |
| MS [%] | 0N | 48.606 ± 1.410 | 63.343 ± 1.250 | 31.128 ± 3.855 | *** <0.001 | 0.991 | 0.494 | w20-22 | 0.992 | 1.000 | *** <0.001 | *** <0.001 | - | - |
| | | | | | *** <0.001 | * 0.028 | *** <0.001 | w22-24 | 0.601 | ** 0.004 | ** 0.006 | *** <0.001 | | |
| | 8N | 64.556 ± 2.224 | 73.002 ± 1.396 | 13.822 ± 3.229 | *** <0.001 | *** <0.001 | *** <0.001 | *** <0.001 | *** <0.001 | *** <0.001 | w20-22 | 1.000 | 0.334 | I** 0.002 |
| | | | | | *** <0.001 | *** <0.001 | *** <0.001 | ** 0.006 | 0.557 | *** <0.001 | w22-24 | 0.274 | | |
| ES [%] | 0N | 30.173 ± 1.288 | 19.512 ± 0.969 | -34.805 ± 3.436 | 0.569 | 0.054 | 0.999 | w20-22 | 0.100 | 0.950 | *** <0.001 | *** <0.001 | - | - |
| | | | | | *** <0.001 | 0.717 | *** <0.001 | w22-24 | 0.077 | * 0.015 | * 0.033 | *** <0.001 | | |
| | 8N | 18.071 ± 1.458 | 13.395 ± 0.637 | -22.858 ± 5.636 | *** <0.001 | ** 0.005 | *** <0.001 | *** <0.001 | 0.102 | *** <0.001 | w20-22 | 0.383 | S* 0.028 | 0.620 |
| | | | | | *** <0.001 | *** <0.001 | *** <0.001 | * 0.033 | 1.000 | *** <0.001 | w22-24 | 0.053 | | |

| | | | | | | | | | | | | | | |
|---|---|---|---|---|---|---|---|---|---|---|---|---|---|---|
| **SclAB (0N: n=9; 8N: n=8)** | | | | | | | | | | | | | | |
| BFR [%/d] | 0N | 1.462 ± 0.136 | 1.639 ± 0.109 | 20.318 ± 15.922 | 0.467 | 0.998 | 1.000 | 0.984 | w20-22 | 0.952 | ** 0.003 | *** <0.001 | - | - |
| | | | | | 0.256 | 0.536 | ** 0.006 | 0.997 | w22-24 | 0.990 | 0.999 | 0.731 | | |
| | 8N | 2.477 ± 0.351 | 1.928 ± 0.150 | -16.789 ± 6.327 | *** <0.001 | *** <0.001 | *** <0.001 | * 0.013 | *** <0.001 | * 0.033 | 1.000 | w20-22 | 0.525 | 0.594 |
| | | | | | ** 0.001 | ** 0.006 | *** <0.001 | 0.291 | 0.731 | 0.227 | 0.379 | w22-24 | | |
| BRR [%/d] | 0N | 1.211 ± 0.081 | 0.474 ± 0.069 | -60.942 ± 4.569 | *** <0.001 | 0.997 | 0.800 | 0.547 | w20-22 | 0.147 | * 0.021 | ** 0.009 | - | - |
| | | | | | *** <0.001 | 0.100 | 0.962 | 0.999 | w22-24 | 0.981 | 0.576 | 0.155 | | |
| | 8N | 0.545 ± 0.061 | 0.174 ± 0.034 | -67.050 ± 6.237 | *** <0.001 | ** 0.007 | 0.346 | *** <0.001 | ** 0.009 | 0.954 | 1.000 | w20-22 | 0.069 | **S* 0.028** |
| | | | | | *** <0.001 | *** <0.001 | ** 0.009 | * 0.028 | 0.155 | * 0.013 | 0.991 | w22-24 | | |
| MAR [µm/d] | 0N | 1.242 ± 0.024 | 1.166 ± 0.031 | -5.981 ± 2.605 | 0.952 | 0.285 | 0.940 | 0.437 | w20-22 | * 0.039 | * 0.012 | * 0.015 | - | - |
| | | | | | 0.403 | 0.869 | 1.000 | 1.000 | w22-24 | 0.130 | 1.000 | 0.787 | | |
| | 8N | 1.499 ± 0.090 | 1.260 ± 0.063 | -15.104 ± 3.318 | * 0.046 | 0.500 | 0.264 | 0.761 | * 0.015 | 1.000 | 1.000 | w20-22 | **S* 0.047** | 0.070 |
| | | | | | 1.000 | 0.999 | 0.643 | 0.698 | 0.787 | 0.955 | 0.922 | w22-24 | | |
| MRR [µm/d] | 0N | 1.968 ± 0.031 | 1.694 ± 0.075 | -13.971 ± 3.383 | 0.664 | 0.986 | *** <0.001 | 0.993 | w20-22 | *** <0.001 | 0.080 | 0.382 | - | - |
| | | | | | 1.000 | 1.000 | *** <0.001 | 0.989 | w22-24 | *** <0.001 | 0.957 | 1.000 | | |
| | 8N | 1.778 ± 0.049 | 1.703 ± 0.093 | -4.080 ± 5.144 | ** 0.001 | 0.656 | *** <0.001 | 0.068 | 0.382 | *** <0.001 | 0.998 | w20-22 | 0.845 | 0.970 |
| | | | | | 1.000 | 1.000 | *** <0.001 | 0.984 | 1.000 | *** <0.001 | 0.945 | w22-24 | | |
| MS [%] | 0N | 50.789 ± 2.387 | 68.026 ± 1.861 | 35.700 ± 5.946 | *** <0.001 | 1.000 | 0.128 | 0.992 | w20-22 | 0.999 | *** <0.001 | *** <0.001 | - | - |
| | | | | | *** <0.001 | *** <0.001 | *** <0.001 | 0.601 | w22-24 | *** <0.001 | 0.557 | ** 0.001 | | |
| | 8N | 65.647 ± 1.951 | 79.382 ± 1.799 | 21.229 ± 2.148 | *** <0.001 | *** <0.001 | *** <0.001 | *** <0.001 | *** <0.001 | *** <0.001 | 1.000 | w20-22 | 0.222 | **I** 0.005** |
| | | | | | *** <0.001 | *** <0.001 | *** <0.001 | *** <0.001 | ** 0.001 | *** <0.001 | 0.274 | w22-24 | | |
| ES [%] | 0N | 24.190 ± 1.589 | 13.979 ± 1.099 | -41.486 ± 3.476 | *** <0.001 | 1.000 | * 0.024 | 0.100 | w20-22 | 0.711 | 0.102 | *** <0.001 | - | - |
| | | | | | *** <0.001 | *** <0.001 | *** <0.001 | 0.077 | w22-24 | *** <0.001 | 1.000 | * 0.023 | | |
| | 8N | 13.231 ± 1.197 | 7.259 ± 0.929 | -43.256 ± 8.665 | *** <0.001 | *** <0.001 | *** <0.001 | *** <0.001 | *** <0.001 | *** <0.001 | 0.383 | w20-22 | **S** 0.005** | **S*** <0.001** |
| | | | | | *** <0.001 | *** <0.001 | *** <0.001 | *** <0.001 | * 0.023 | *** <0.001 | 0.053 | w22-24 | | |

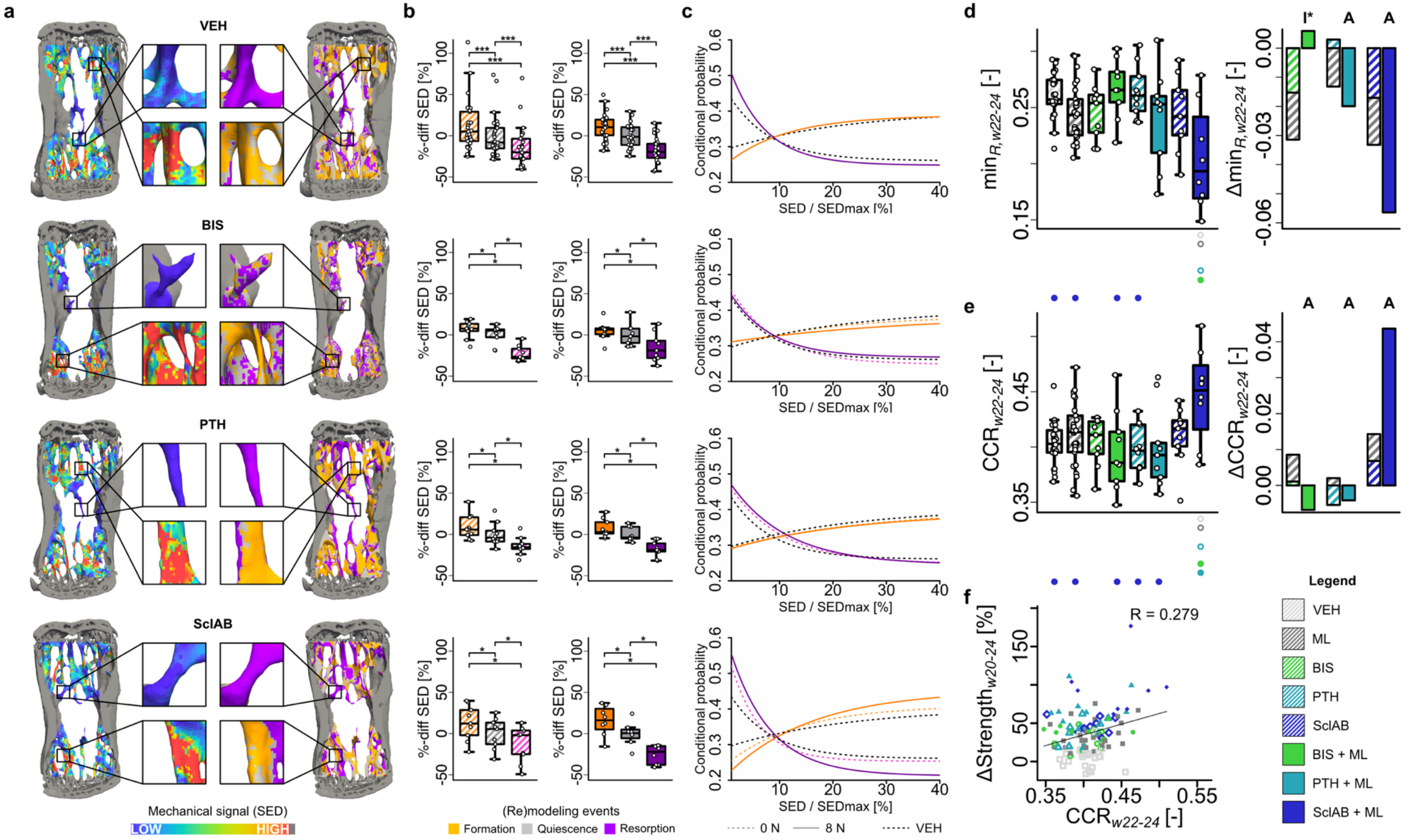


**Figure S.3: Formation aligns with high mechanical signal, resorption with low: (a)** Spatial comparison of local mechanical signal (SED at week 20) with sites of trabecular bone formation and resorption (w22-24). **(b)** SED at formation, resorption, and quiescent sites[2], normalized to the group mean of quiescence to allow statistical testing with this group. Each data point represents one animal (biological replicate; VEH: n=25, ML: n=26, BIS 0N: n=9, BIS 8N: n=9, PTH 0N: n=10, PTH 8N: n=9, SclAB 0N: n=9, SclAB 8N: n=8). The box spans the interquartile range (IQR) from the 25th to the 75th percentile. The median is shown as a line inside the box. Whiskers extend to the most extreme data points within 1.5 × IQR, points beyond this range are plotted as outliers. *p<0.05, ***<0.001 using two-sided paired Wilcoxon-signed rank test with Bonferroni correction; exact p-values are found in Supplementary Table S.6. **(c)** The plots show the exponential fitting functions (taking all animals per group into account) for conditional probability of formation and resorption as a function of SED/SEDmax. Dashed lines represent monotreatment (0N, including VEH), and solid lines represent combined treatment with mechanical loading (8N, including ML). VEH is indicated by black dotted line. Values are cropped above 40% due to low voxel count[2]. **(d-e)** Each data point represents one animal (biological replicate; VEH: n=25, ML: n=26, BIS 0N: n=9, BIS 8N: n=9, PTH 0N: n=10, PTH 8N: n=9, SclAB 0N: n=9, SclAB 8N: n=8). The box spans the interquartile range (IQR) from the 25th to the 75th percentile. The median is shown as a line inside the box. Whiskers extend to the most extreme data points within 1.5 × IQR, points beyond this range are plotted as outliers. [-] denotes unitless, Δ denotes change from VEH; sum of single vs. combined treatment group means indicating antagonistic (I), additive (A) or synergistic (S) effects. **(d)** Asymptotic resorption probability value) **(e)** CCR as a measure of mechanoregulated (re)modeling **(f)** Linear regression analysis in week interval w22-24 of ΔStrength with CCR. Significant differences are depicted with a symbol in the color of the corresponding group (o: to monotreatment, ●: to combined treatment, see Supplementary Table S.1 for color encoding, p<0.05). Source data are provided as a Source Data file.

**Table S.6:** Group means ± standard error of the absolute strain energy density (SED) values in formation (F), quiescence (Q) and resorption (R) sites are reported for all vehicle and treatment groups for both time intervals. Furthermore, mean SED at formation, quiescent and resorption sites expressed as percent difference from the mean (over all animals) quiescent SED are reported as |F-Q| and |Q-R|. Normalization to the mean (over all animals) quiescent SED preserves group properties thus allows statistical comparison of relative differences between F,Q,R regions. P-values were calculated using pairwise two-sided Wilcoxon signed-rank tests with Bonferroni correction. Significance is indicated as: *p<0.05, **p<0.01, ***p<0.001.

| Group | Week | F [kPa] | Q [kPa] | R [kPa] | \|F-Q\| [%] | \|Q-R\| [%] | p-value (F vs. Q) | p-value (Q vs. R) |
|---|---|---|---|---|---|---|---|---|
| VEH 0N | 20-22 | 3.219 ± 0.270 | 2.882 ± 0.216 | 2.453 ± 0.157 | -9.466 ± 1.079 | -13.912 ± 1.149 | ***<br>**<0.001** | ***<br>**<0.001** |
| | 22-24 | 3.182 ± 0.178 | 2.820 ± 0.146 | 2.430 ± 0.140 | -10.818 ± 1.105 | -14.064 ± 1.245 | ***<br>**<0.001** | ***<br>**<0.001** |
| VEH 8N | 20-22 | 11.007 ± 0.474 | 9.575 ± 0.362 | 7.649 ± 0.349 | -12.358 ± 1.499 | -20.396 ± 1.563 | ***<br>**<0.001** | ***<br>**<0.001** |
| | 22-24 | 8.184 ± 0.240 | 7.433 ± 0.211 | 6.107 ± 0.224 | -9.008 ± 0.944 | -17.932 ± 1.761 | ***<br>**<0.001** | ***<br>**<0.001** |
| BIS 0N | 20-22 | 2.962 ± 0.203 | 2.765 ± 0.176 | 2.269 ± 0.143 | -6.275 ± 1.429 | -17.742 ± 2.013 | *<br>**0.012** | *<br>**0.012** |
| | 22-24 | 2.282 ± 0.077 | 2.143 ± 0.082 | 1.663 ± 0.066 | -6.176 ± 1.235 | -21.962 ± 2.789 | *<br>**0.023** | *<br>**0.012** |
| BIS 8N | 20-22 | 10.552 ± 0.469 | 9.836 ± 0.433 | 8.268 ± 0.552 | -6.640 ± 1.729 | -16.333 ± 3.029 | *<br>**0.023** | *<br>**0.012** |
| | 22-24 | 7.409 ± 0.265 | 7.066 ± 0.311 | 5.998 ± 0.407 | -4.542 ± 2.560 | -15.696 ± 2.612 | n.s.<br>0.750 | *<br>**0.012** |
| PTH 0N | 20-22 | 3.142 ± 0.257 | 2.841 ± 0.210 | 2.291 ± 0.186 | -9.022 ± 1.643 | -19.408 ± 1.848 | *<br>**0.012** | **<br>**0.006** |
| | 22-24 | 2.353 ± 0.100 | 2.162 ± 0.092 | 1.854 ± 0.071 | -8.051 ± 1.232 | -14.092 ± 1.161 | **<br>**0.006** | **<br>**0.006** |

| | | | | | | | | |
|---|---|---|---|---|---|---|---|---|
| PTH 8N | 20-22 | 10.192 ± 0.672 | 8.341 ± 0.481 | 6.661 ± 0.501 | -17.642 ± 2.301 | -20.566 ± 1.610 | * **0.012** | * **0.012** |
| | 22-24 | 5.835 ± 0.199 | 5.435 ± 0.157 | 4.469 ± 0.175 | -6.673 ± 1.225 | -17.568 ± 3.207 | * **0.012** | * **0.023** |
| SclAB 0N | 20-22 | 3.185 ± 0.213 | 2.654 ± 0.172 | 2.246 ± 0.145 | -16.350 ± 2.545 | -15.293 ± 1.466 | * **0.012** | * **0.012** |
| | 22-24 | 2.333 ± 0.148 | 2.069 ± 0.128 | 1.799 ± 0.152 | -11.208 ± 0.866 | -13.966 ± 3.199 | * **0.012** | * **0.023** |
| SclAB 8N | 20-22 | 13.647 ± 1.477 | 11.073 ± 1.173 | 8.818 ± 1.399 | -18.578 ± 1.754 | -22.010 ± 4.365 | * **0.023** | * **0.023** |
| | 22-24 | 6.971 ± 0.380 | 6.032 ± 0.296 | 4.487 ± 0.248 | -12.978 ± 2.650 | -25.266 ± 3.306 | * **0.023** | * **0.023** |

**Table S.7:** Saturating exponential curve fits for formation slope values were conducted by fitting the function $f(x) = y_0 + a \cdot (1 - \exp(-b \cdot x))$ to the first 40% of the normalized SED/SED$_{max}$ interval for each animal individually. Coefficients are presented as group means ± standard error. Parameter $y_0$ reflects the level of formation in the absence of mechanical stimulus (i.e. at SED = 0), representing non-targeted formation. Coefficient $a$ indicates the maximal increase in formation in response to SED, while $y_0 + a$ defines the plateau, or the maximum potential formation probability at high SED. $b$ represents the sensitivity or rate of response, i.e. higher $b$-values indicate that the curve rises more quickly, reaching the plateau earlier (i.e. formation increases rapidly with even low SED). Synergy (S) or antagonism (I) between treatment and mechanical loading was assessed using the interaction term “treatment:loading” from a fitted linear model applied to the log$_2$-transformed absolute values within each interval. P-values were obtained using one-way ANOVA in each time interval for the shown groups followed by Tukey’s post-hoc test to account for multiple comparisons. ‘w20-22’ or ’w22-24’ denotes the reference group; asterisks (*, **, ***) indicate groups that are significantly different from it. Significance is indicated as: *p<0.05, **p<0.01, ***p<0.001.

| **Readout** | **N** | **w20-22** | **w22-24** | **%-change between w20-22 and w22-24 [%]** | ○ | ○ | ○ | ○ | ○ | ● | ● | ● | Synergy w20-22 | Synergy w22-24 |
|---|---|---|---|---|---|---|---|---|---|---|---|---|---|---|
| **VEH (0N: n=25; 8N: n=26)** | | | | | | | | | | | | | | |
| y0 (level of formation at SED=0) | 0N | 0.297 ± 0.004 | 0.291 ± 0.005 | -1.974 ± 1.475 | w20-22 | * **0.020** | 0.997 | 0.721 | * **0.014** | 0.999 | *** **<0.001** | *** **<0.001** | - | - |
| | | | | | w22-24 | ** **0.003** | 1.000 | 0.920 | ** **0.002** | 0.991 | * **0.029** | *** **<0.001** | | |
| | 8N | 0.269 ± 0.007 | 0.251 ± 0.009 | -6.011 ± 2.847 | * **0.020** | w20-22 | * **0.042** | 0.984 | 0.959 | 0.056 | 0.518 | 0.157 | - | - |
| | | | | | ** **0.003** | w22-24 | 0.051 | 0.637 | 0.907 | ** **0.007** | 1.000 | 0.181 | | |
| a (maximum increase in formation) | 0N | 0.170 ± 0.036 | 0.182 ± 0.031 | -9.563 ± 73.721 | w20-22 | 0.774 | 0.998 | 1.000 | 0.996 | 0.838 | ** **0.005** | 0.965 | - | - |
| | | | | | w22-24 | 1.000 | 1.000 | 1.000 | 0.977 | 1.000 | 0.767 | 0.302 | | |
| | 8N | 0.251 ± 0.038 | 0.166 ± 0.022 | 4.991 ± 17.610 | 0.774 | w20-22 | 1.000 | 0.748 | 1.000 | 0.179 | 0.119 | 1.000 | - | - |
| | | | | | 1.000 | w22-24 | 1.000 | 1.000 | 0.921 | 1.000 | 0.605 | 0.187 | | |
| b (sensitivity of response) | 0N | 0.035 ± 0.006 | 0.044 ± 0.007 | 182.258 ± 85.085 | w20-22 | 0.889 | 1.000 | 0.956 | 0.971 | 1.000 | 0.992 | 0.276 | - | - |
| | | | | | w22-24 | * **0.028** | 1.000 | 1.000 | 0.172 | 0.999 | 0.887 | 0.997 | | |
| | 8N | 0.048 ± 0.008 | 0.084 ± 0.009 | 378.359 ± 131.954 | 0.889 | w20-22 | 0.955 | 1.000 | 1.000 | 0.898 | 1.000 | 0.831 | - | - |

| | | | | | | | | | | | | | | |
|---|---|---|---|---|---|---|---|---|---|---|---|---|---|---|
| | | | | | * 0.028 | w22-24 | 0.221 | 0.299 | 1.000 | 0.076 | 0.963 | 0.760 | | |
| y0+a (plateau at high mechanical strains) | 0N | 0.467 ± 0.035 | 0.473 ± 0.031 | 8.536 ± 7.584 | w20-22 | 0.970 | 0.994 | 0.997 | 1.000 | 0.874 | * 0.039 | 1.000 | - | - |
| | | | | | w22-24 | 0.942 | 1.000 | 1.000 | 1.000 | 1.000 | 0.984 | 0.918 | | |
| | 8N | 0.520 ± 0.038 | 0.417 ± 0.020 | -11.339 ± 6.734 | 0.970 | w20-22 | 1.000 | 0.821 | 0.999 | 0.421 | 0.222 | 1.000 | - | - |
| | | | | | 0.942 | w22-24 | 0.985 | 0.998 | 0.976 | 0.999 | 0.652 | 0.457 | | |
| **BIS (0N: n=9; 8N: n=9)** | | | | | | | | | | | | | | |
| y0 | 0N | 0.305 ± 0.003 | 0.293 ± 0.007 | -3.918 ± 1.831 | 0.997 | * 0.042 | w20-22 | 0.540 | * 0.016 | 1.000 | ** 0.001 | *** <0.001 | - | - |
| | | | | | 1.000 | 0.051 | w22-24 | 0.940 | * 0.012 | 0.999 | 0.090 | *** <0.001 | | |
| | 8N | 0.304 ± 0.010 | 0.302 ± 0.011 | -0.043 ± 4.188 | 0.999 | 0.056 | 1.000 | 0.601 | * 0.021 | w20-22 | ** 0.002 | *** <0.001 | 0.124 | **I* 0.020** |
| | | | | | 0.991 | ** 0.007 | 0.999 | 0.682 | ** 0.002 | w22-24 | * 0.021 | *** <0.001 | | |
| a | 0N | 0.218 ± 0.065 | 0.184 ± 0.061 | 77.465 ± 69.861 | 0.998 | 1.000 | w20-22 | 0.984 | 1.000 | 0.665 | 0.156 | 1.000 | - | - |
| | | | | | 1.000 | 1.000 | w22-24 | 1.000 | 0.994 | 1.000 | 0.908 | 0.553 | | |
| | 8N | 0.066 ± 0.056 | 0.151 ± 0.047 | -326.699 ± 212.075 | 0.838 | 0.179 | 0.665 | 0.988 | 0.640 | w20-22 | *** <0.001 | 0.468 | 0.206 | 0.683 |
| | | | | | 1.000 | 1.000 | 1.000 | 1.000 | 0.932 | w22-24 | 0.699 | 0.303 | | |
| b | 0N | 0.033 ± 0.009 | 0.042 ± 0.014 | 658.509 ± 610.023 | 1.000 | 0.955 | w20-22 | 0.971 | 0.980 | 1.000 | 0.993 | 0.433 | - | - |
| | | | | | 1.000 | 0.221 | w22-24 | 1.000 | 0.362 | 1.000 | 0.942 | 0.998 | | |
| | 8N | 0.030 ± 0.011 | 0.035 ± 0.013 | 164.000 ± 138.631 | 1.000 | 0.898 | 1.000 | 0.936 | 0.952 | w20-22 | 0.980 | 0.339 | 0.444 | **S* 0.015** |
| | | | | | 0.999 | 0.076 | 1.000 | 0.999 | 0.175 | w22-24 | 0.785 | 0.970 | | |
| y0+a | 0N | 0.523 ± 0.067 | 0.478 ± 0.061 | 5.029 ± 19.776 | 0.994 | 1.000 | w20-22 | 0.920 | 0.999 | 0.644 | 0.498 | 1.000 | - | - |
| | | | | | 1.000 | 0.985 | w22-24 | 1.000 | 1.000 | 1.000 | 0.997 | 0.976 | | |
| | 8N | 0.370 ± 0.049 | 0.453 ± 0.047 | 145.241 ± 137.128 | 0.874 | 0.421 | 0.644 | 0.999 | 0.918 | w20-22 | ** 0.007 | 0.897 | **I** 0.007** | 0.682 |
| | | | | | 1.000 | 0.999 | 1.000 | 1.000 | 1.000 | w22-24 | 0.973 | 0.905 | | |
| **PTH (0N: n=10; 8N: n=9)** | | | | | | | | | | | | | | |
| y0 | 0N | 0.279 ± 0.007 | 0.275 ± 0.008 | -1.031 ± 3.629 | 0.721 | 0.984 | 0.540 | w20-22 | 0.716 | 0.601 | 0.260 | 0.701 | - | - |
| | | | | | 0.920 | 0.637 | 0.940 | w22-24 | 0.204 | 0.682 | 0.636 | * 0.012 | | |
| | 8N | 0.246 ± 0.009 | 0.246 ± 0.010 | 0.629 ± 4.501 | *** 0.001 | 0.518 | ** 0.001 | 0.260 | 0.996 | ** 0.002 | w20-22 | 0.998 | 0.600 | 0.631 |
| | | | | | * 0.029 | 1.000 | 0.090 | 0.636 | 0.996 | * 0.021 | w22-24 | 0.612 | | |
| a | 0N | 0.140 ± 0.022 | 0.185 ± 0.043 | 41.451 ± 27.850 | 1.000 | 0.748 | 0.984 | w20-22 | 0.980 | 0.988 | * 0.011 | 0.918 | - | - |
| | | | | | 1.000 | 1.000 | 1.000 | w22-24 | 0.993 | 1.000 | 0.897 | 0.524 | | |
| | 8N | 0.449 ± 0.111 | 0.288 ± 0.101 | -2.790 ± 38.514 | ** 0.005 | 0.119 | 0.156 | * 0.011 | 0.169 | *** <0.001 | w20-22 | 0.354 | 0.119 | 0.360 |
| | | | | | 0.767 | 0.605 | 0.908 | 0.897 | 1.000 | 0.699 | w22-24 | 0.998 | | |
| b | 0N | 0.050 ± 0.008 | 0.046 ± 0.010 | -3.976 ± 23.171 | 0.956 | 1.000 | 0.971 | w20-22 | 1.000 | 0.936 | 1.000 | 0.944 | - | - |

| | | | | | | | | | | | | | | |
|---|---|---|---|---|---|---|---|---|---|---|---|---|---|---|
| | | | | | 1.000 | 0.299 | 1.000 | w22-24 | 0.455 | 0.999 | 0.977 | 1.000 | | |
| | 8N | 0.046 ± 0.016 | 0.066 ± 0.013 | 456.616 ± 263.598 | 0.992 | 1.000 | 0.993 | 1.000 | 1.000 | 0.980 | w20-22 | 0.891 | 0.152 | 0.399 |
| | | | | | 0.887 | 0.963 | 0.942 | 0.977 | 0.966 | 0.785 | w22-24 | 1.000 | | |
| y0+a | 0N | 0.419 ± 0.020 | 0.459 ± 0.043 | 8.605 ± 6.560 | 0.997 | 0.821 | 0.920 | w20-22 | 0.997 | 0.999 | * 0.031 | 0.994 | - | - |
| | | | | | 1.000 | 0.998 | 1.000 | w22-24 | 1.000 | 1.000 | 0.981 | 0.921 | | |
| | 8N | 0.695 ± 0.115 | 0.534 ± 0.100 | -9.314 ± 19.146 | * 0.039 | 0.222 | 0.498 | * 0.031 | 0.204 | ** 0.007 | w20-22 | 0.285 | 0.101 | 0.220 |
| | | | | | 0.984 | 0.652 | 0.997 | 0.981 | 0.998 | 0.973 | w22-24 | 1.000 | | |
| **SclAB (0N: n=9; 8N: n=8)** | | | | | | | | | | | | | | |
| y0 | 0N | 0.256 ± 0.012 | 0.234 ± 0.015 | -8.253 ± 4.550 | * 0.014 | 0.959 | * 0.016 | 0.716 | w20-22 | * 0.021 | 0.996 | 0.886 | - | - |
| | | | | | ** 0.002 | 0.907 | * 0.012 | 0.204 | w22-24 | ** 0.002 | 0.996 | 0.950 | | |
| | 8N | 0.237 ± 0.016 | 0.215 ± 0.018 | -6.395 ± 10.024 | *** <0.001 | 0.157 | *** <0.001 | 0.071 | 0.886 | *** <0.001 | 0.998 | w20-22 | 0.819 | 0.450 |
| | | | | | *** <0.001 | 0.181 | *** <0.001 | * 0.012 | 0.950 | *** <0.001 | 0.6122 | w22-24 | | |
| a | 0N | 0.221 ± 0.036 | 0.248 ± 0.067 | 22.782 ± 32.467 | 0.996 | 1.000 | 1.000 | 0.980 | w20-22 | 0.640 | 0.169 | 1.000 | - | - |
| | | | | | 0.977 | 0.921 | 0.994 | 0.993 | w22-24 | 0.932 | 1.000 | 0.945 | | |
| | 8N | 0.249 ± 0.038 | 0.345 ± 0.081 | 54.781 ± 35.118 | 0.965 | 1.000 | 1.000 | 0.918 | 1.000 | 0.468 | 0.354 | w20-22 | 0.470 | 0.258 |
| | | | | | 0.302 | 0.187 | 0.553 | 0.524 | 0.945 | 0.303 | 0.998 | w22-24 | | |
| b | 0N | 0.049 ± 0.012 | 0.087 ± 0.022 | 102.185 ± 51.133 | 0.971 | 1.000 | 0.980 | 1.000 | w20-22 | 0.952 | 1.000 | 0.942 | - | - |
| | | | | | 0.172 | 1.000 | 0.362 | 0.455 | w22-24 | 0.175 | 0.966 | 0.814 | | |
| | 8N | 0.069 ± 0.018 | 0.056 ± 0.013 | 47.010 ± 58.956 | 0.276 | 0.831 | 0.433 | 0.944 | 0.942 | 0.339 | 0.891 | w20-22 | 0.882 | **I* 0.032** |
| | | | | | 0.997 | 0.760 | 0.998 | 1.000 | 0.814 | 0.970 | 1.000 | w22-24 | | |
| y0+a | 0N | 0.478 ± 0.033 | 0.483 ± 0.070 | 2.365 ± 13.366 | 1.000 | 0.999 | 0.999 | 0.997 | w20-22 | 0.918 | 0.204 | 1.000 | - | - |
| | | | | | 1.000 | 0.976 | 1.000 | 1.000 | w22-24 | 1.000 | 0.998 | 0.983 | | |
| | 8N | 0.486 ± 0.034 | 0.560 ± 0.083 | 21.002 ± 22.630 | 1.000 | 1.000 | 1.000 | 0.994 | 1.000 | 0.897 | 0.285 | w20-22 | 0.676 | 0.107 |
| | | | | | 0.918 | 0.457 | 0.976 | 0.921 | 0.983 | 0.905 | 1.000 | w22-24 | | |

**Table S.8:** Decaying exponential curve fits for resorption were conducted by fitting the function $f(x) = y_0 + a \cdot \exp(-b \cdot x)$ to the first 40% of the normalized SED/$SED_{max}$ interval for each animal individually. The fitted coefficients for each animal are presented as group means ± standard error. Resorption asymptote $min_R$ represents the saturation level of resorption at high mechanical signal, i.e. the minimum resorption probability ($x \to \infty$), indicative of the amount of non-targeted resorption. Coefficient $a$ represents the magnitude of the response, i.e. the difference between the initial and the minimum resorption probability. $a + y_0$ represents the peak resorption probability at low SED. Coefficient $b$ is the rate of decay, representing how quickly the probability of resorption drops as SED increases. Synergy (S) or antagonism (I) between treatment and mechanical loading was assessed using the interaction term “treatment:loading” from a fitted linear model applied to the $log_2$-transformed absolute values within each interval. P-values were obtained using one-way ANOVA in each time interval for the shown groups followed by Tukey’s post-hoc test to account for multiple comparisons. ‘w20-22’ or ‘w22-24’ denotes the reference group; asterisks (*, **, ***) indicate groups that are significantly different from it. Significance is indicated as: *p<0.05, **p<0.01, ***p<0.001.

| Readout | N | w20-22 | w22-24 | %-change between w20-22 and w22-24 [%] | ○ | ○ | ○ | ○ | ○ | ● | ● | ● | Synergy w20-22 | Synergy w22-24 |
|---|---|---|---|---|---|---|---|---|---|---|---|---|---|---|
| **VEH (0N: n=25; 8N: n=26)** | | | | | | | | | | | | | | |
| minR (saturation level of resorption; plateau at high mechanical strains) | 0N | 0.266 ± 0.003 | 0.261 ± 0.004 | -1.845 ± 1.304 | w20-22 | 0.020 * | 0.977 | 0.997 | 0.740 | 0.549 | 0.002 ** | 0.070 | - | - |
| | | | | | w22-24 | 0.474 | 0.867 | 1.000 | 0.786 | 0.999 | 0.625 | <0.001 *** | | |
| | 8N | 0.233 ± 0.006 | 0.245 ± 0.005 | 7.366 ± 5.030 | 0.020 * | w20-22 | 0.804 | 0.579 | 0.989 | 0.999 | 0.738 | 0.999 | - | - |
| | | | | | 0.474 | w22-24 | 1.000 | 0.622 | 1.000 | 0.490 | 1.000 | 0.016 * | | |
| a (magnitude of response) | 0N | 0.200 ± 0.010 | 0.208 ± 0.011 | 6.873 ± 6.119 | w20-22 | 0.002 ** | 0.999 | 0.596 | 0.067 | 1.000 | 0.006 ** | <0.001 *** | - | - |
| | | | | | w22-24 | <0.001 *** | 0.968 | 0.988 | 0.012 * | 1.000 | 0.029 * | <0.001 *** | | |
| | 8N | 0.274 ± 0.014 | 0.304 ± 0.015 | 15.330 ± 6.368 | 0.002 ** | w20-22 | 0.015 * | 0.928 | 1.000 | 0.037 * | 0.988 | 0.159 | - | - |
| | | | | | <0.001 *** | w22-24 | 0.204 | 0.105 | 1.000 | 0.018 * | 1.000 | 0.013 * | | |
| b (sensitivity of decay) | 0N | 0.144 ± 0.006 | 0.145 ± 0.008 | 6.571 ± 7.975 | w20-22 | 0.910 | 0.712 | 1.000 | 1.000 | 0.407 | 0.332 | 0.491 | - | - |
| | | | | | w22-24 | 0.988 | 0.917 | 0.884 | 0.997 | 1.000 | 0.667 | 1.000 | | |
| | 8N | 0.130 ± 0.008 | 0.157 ± 0.012 | 832.154 ± 820.041 | 0.910 | w20-22 | 0.995 | 1.000 | 0.978 | 0.924 | 0.878 | 0.088 | - | - |
| | | | | | 0.988 | w22-24 | 0.571 | 0.491 | 1.000 | 1.000 | 0.269 | 0.999 | | |
| y0+a (peak resorption under zero mechanical loading) | 0N | 0.467 ± 0.008 | 0.469 ± 0.009 | 0.897 ± 2.038 | w20-22 | 0.126 | 0.896 | 0.639 | 0.174 | 0.783 | 0.495 | <0.001 *** | - | - |
| | | | | | w22-24 | <0.001 *** | 0.999 | 0.898 | 0.005 ** | 1.000 | 0.024 * | <0.001 *** | | |
| | 8N | 0.507 ± 0.013 | 0.548 ± 0.012 | 9.476 ± 3.470 | 0.126 | w20-22 | 0.030 * | 1.000 | 0.998 | 0.014 * | 1.000 | 0.110 | - | - |
| | | | | | <0.001 *** | w22-24 | 0.033 * | 0.130 | 1.000 | 0.018 * | 1.000 | 0.107 | | |
| **BIS (0N: n=9; 8N: n=9)** | | | | | | | | | | | | | | |
| minR | 0N | 0.253 ± 0.009 | 0.246 ± 0.008 | -2.098 ± 4.855 | 0.977 | 0.804 | w20-22 | 1.000 | 0.999 | 0.994 | 0.183 | 0.682 | - | - |

| | | | | | | | | | | | | | | |
|---|---|---|---|---|---|---|---|---|---|---|---|---|---|---|
| | | | | | 0.867 | 1.000 | w22-24 | 0.860 | 1.000 | 0.766 | 1.000 | 0.071 | | |
| | 8N | 0.241 ± 0.022 | 0.267 ± 0.009 | 26.748 ± 24.035 | 0.549 | 0.999 | 0.994 | 0.968 | 1.000 | w20-22 | 0.629 | 0.979 | 0.692 | **I* 0.027** |
| | | | | | 0.999 | 0.490 | 0.766 | 1.000 | 0.684 | w22-24 | 0.542 | *** <0.001 | | |
| a | 0N | 0.186 ± 0.016 | 0.236 ± 0.018 | 30.103 ± 9.802 | 0.999 | * 0.015 | w20-22 | 0.505 | 0.078 | 1.000 | * 0.012 | *** <0.001 | - | - |
| | | | | | 0.968 | 0.204 | w22-24 | 1.000 | 0.421 | 0.994 | 0.577 | *** <0.001 | | |
| | 8N | 0.194 ± 0.031 | 0.211 ± 0.022 | 27.676 ± 17.435 | 1.000 | * 0.037 | 1.000 | 0.673 | 0.141 | w20-22 | * 0.025 | *** <0.001 | **I* 0.024** | **I** 0.004** |
| | | | | | 1.000 | * 0.018 | 0.994 | 0.998 | 0.088 | w22-24 | 0.154 | *** <0.001 | | |
| b | 0N | 0.119 ± 0.014 | 0.121 ± 0.008 | 22.051 ± 22.906 | 0.712 | 0.995 | w20-22 | 0.965 | 0.848 | 1.000 | 1.000 | 0.065 | - | - |
| | | | | | 0.917 | 0.571 | w22-24 | 1.000 | 0.761 | 0.864 | 1.000 | 0.975 | | |
| | 8N | 0.112 ± 0.016 | 0.153 ± 0.022 | 156.850 ± 136.350 | 0.407 | 0.924 | 1.000 | 0.834 | 0.624 | w20-22 | 1.000 | * 0.023 | 0.807 | 0.589 |
| | | | | | 1.000 | 1.000 | 0.864 | 0.830 | 1.000 | w22-24 | 0.632 | 1.000 | | |
| y0+a | 0N | 0.440 ± 0.013 | 0.482 ± 0.010 | 10.034 ± 2.672 | 0.896 | * 0.030 | w20-22 | 0.189 | * 0.037 | 1.000 | 0.131 | *** <0.001 | - | - |
| | | | | | | * | w22-24 | | | | | *** | | |
| | 8N | 0.435 ± 0.019 | 0.477 ± 0.017 | 10.423 ± 2.529 | 0.783 | * 0.014 | 1.000 | 0.121 | * 0.021 | w20-22 | 0.082 | *** <0.001 | 0.127 | **I** 0.004** |
| | | | | | 1.000 | * 0.018 | 1.000 | 0.996 | 0.093 | w22-24 | 0.237 | *** <0.001 | | |
| **PTH (0N: n=10; 8N: n=9)** | | | | | | | | | | | | | | |
| $min_R$ | 0N | 0.257 ± 0.006 | 0.264 ± 0.007 | 3.684 ± 5.304 | 0.997 | 0.579 | 1.000 | w20-22 | 0.993 | 0.968 | 0.090 | 0.497 | - | - |
| | | | | | 1.000 | 0.622 | 0.860 | w22-24 | 0.792 | 1.000 | 0.658 | *** <0.001 | | |
| | 8N | 0.212 ± 0.019 | 0.241 ± 0.014 | 34.784 ± 30.913 | ** 0.002 | 0.738 | 0.183 | 0.090 | 0.468 | 0.629 | w20-22 | 0.994 | 0.277 | 0.578 |
| | | | | | 0.625 | 1.000 | 1.000 | 0.658 | 1.000 | 0.542 | w22-24 | 0.159 | | |
| a | 0N | 0.245 ± 0.014 | 0.231 ± 0.016 | -0.844 ± 11.363 | 0.596 | 0.928 | 0.505 | w20-22 | 0.969 | 0.673 | 0.682 | * 0.039 | - | - |
| | | | | | 0.988 | 0.105 | 1.000 | w22-24 | 0.294 | 0.998 | 0.437 | *** <0.001 | | |
| | 8N | 0.296 ± 0.017 | 0.297 ± 0.027 | 0.065 ± 8.468 | ** 0.006 | 0.988 | * 0.012 | 0.682 | 0.998 | * 0.025 | w20-22 | 0.809 | 0.490 | 0.381 |
| | | | | | * 0.029 | 1.000 | 0.577 | 0.437 | 1.000 | 0.154 | w22-24 | * 0.043 | | |
| b | 0N | 0.137 ± 0.012 | 0.121 ± 0.007 | -2.950 ± 14.099 | 1.000 | 1.000 | 0.965 | w20-22 | 1.000 | 0.834 | 0.778 | 0.454 | - | - |
| | | | | | 0.884 | 0.491 | 1.000 | w22-24 | 0.714 | 0.830 | 1.000 | 0.966 | | |
| | 8N | 0.110 ± 0.017 | 0.112 ± 0.010 | 43.069 ± 41.462 | 0.332 | 0.878 | 1.000 | 0.778 | 0.554 | 1.000 | w20-22 | * 0.017 | 0.765 | 0.326 |
| | | | | | 0.667 | 0.269 | 1.000 | 1.000 | 0.498 | 0.632 | w22-24 | 0.864 | | |
| y0+a | 0N | 0.502 ± 0.013 | 0.495 ± 0.011 | -0.762 ± 3.604 | 0.639 | 1.000 | 0.189 | w20-22 | 0.995 | 0.121 | 1.000 | 0.172 | - | - |
| | | | | | 0.898 | 0.130 | 0.999 | w22-24 | 0.348 | 0.996 | 0.630 | *** <0.001 | | |
| | 8N | 0.508 ± 0.017 | 0.538 ± 0.017 | 6.339 ± 3.056 | 0.495 | 1.000 | 0.131 | 1.000 | 1.000 | 0.082 | w20-22 | 0.302 | 0.273 | 0.218 |
| | | | | | * 0.024 | 1.000 | 0.325 | 0.630 | 1.000 | 0.237 | w22-24 | 0.128 | | |

**SclAB (0N: n=9; 8N: n=8)**

| | | | | | | | | | | | | | | |
|---|---|---|---|---|---|---|---|---|---|---|---|---|---|---|
| $min_R$ | 0N | 0.245 ± 0.005 | 0.244 ± 0.011 | -0.269 ± 4.322 | 0.740 | 0.989 | 0.999 | 0.993 | w20-22 | 1.000 | 0.468 | 0.932 | - | - |
| | | | | | 0.786 | 1.000 | 1.000 | 0.792 | w22-24 | 0.684 | 1.000 | 0.099 | | |
| | 8N | 0.225 ± 0.014 | 0.204 ± 0.016 | -7.877 ± 6.875 | 0.070 | 0.999 | 0.682 | 0.497 | 0.932 | 0.979 | 0.994 | w20-22 | 0.717 | 0.074 |
| | | | | | *** <0.001 | * 0.016 | 0.071 | *** <0.001 | 0.099 | *** <0.001 | 0.159 | w22-24 | | |
| a | 0N | 0.276 ± 0.018 | 0.305 ± 0.034 | 8.845 ± 8.983 | 0.067 | 1.000 | 0.078 | 0.969 | w20-22 | 0.141 | 0.998 | 0.400 | - | - |
| | | | | | * 0.012 | 1.000 | 0.421 | 0.294 | w22-24 | 0.088 | 1.000 | 0.079 | | |
| | 8N | 0.343 ± 0.036 | 0.404 ± 0.021 | 29.009 ± 17.894 | *** <0.001 | 0.159 | *** <0.001 | * 0.039 | 0.400 | *** <0.001 | 0.809 | w20-22 | 0.512 | 0.839 |
| | | | | | *** <0.001 | * 0.013 | *** <0.001 | *** <0.001 | 0.079 | *** <0.001 | * 0.043 | w22-24 | | |
| b | 0N | 0.144 ± 0.008 | 0.158 ± 0.015 | 10.653 ± 10.903 | 1.000 | 0.978 | 0.848 | 1.000 | w20-22 | 0.624 | 0.554 | 0.734 | - | - |
| | | | | | 0.997 | 1.000 | 0.761 | 0.714 | w22-24 | 1.000 | 0.498 | 0.999 | | |
| | 8N | 0.175 ± 0.011 | 0.145 ± 0.019 | -14.499 ± 11.321 | 0.491 | 0.088 | 0.065 | 0.454 | 0.734 | * 0.023 | * 0.017 | w20-22 | 0.184 | 0.311 |
| | | | | | 1.000 | 0.999 | 0.975 | 0.966 | 0.999 | 1.000 | 0.864 | w22-24 | | |
| y0+a | 0N | 0.520 ± 0.015 | 0.549 ± 0.027 | 5.211 ± 3.311 | 0.174 | 0.998 | * 0.037 | 0.995 | w20-22 | * 0.021 | 1.000 | 0.604 | - | - |
| | | | | | ** 0.005 | 1.000 | 0.139 | 0.348 | w22-24 | 0.093 | 1.000 | 0.299 | | |
| | 8N | 0.568 ± 0.026 | 0.609 ± 0.014 | 9.076 ± 6.230 | *** <0.001 | 0.110 | *** <0.001 | 0.172 | 0.604 | *** <0.001 | 0.302 | w20-22 | 0.854 | 0.502 |
| | | | | | *** <0.001 | 0.107 | *** <0.001 | *** <0.001 | 0.299 | *** <0.001 | 0.128 | w22-24 | | |

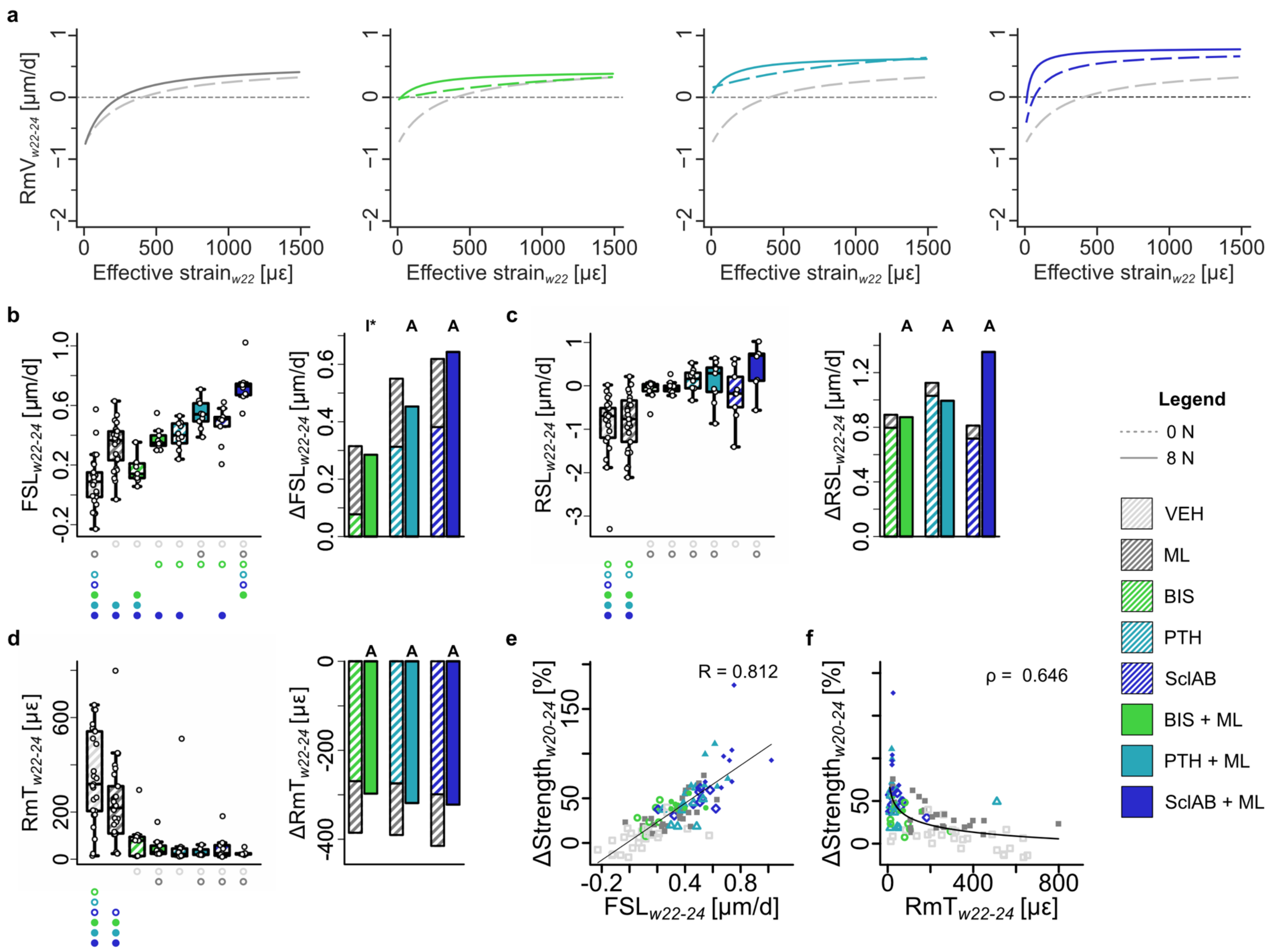
a
b
c
d
e
f
RmV$_{w22-24}$ [µm/d]
Effective strain$_{w22}$ [µε]
FSL$_{w22-24}$ [µm/d]
ΔFSL$_{w22-24}$ [µm/d]
RSL$_{w22-24}$ [µm/d]
ΔRSL$_{w22-24}$ [µm/d]
RmT$_{w22-24}$ [µε]
ΔRmT$_{w22-24}$ [µε]
ΔStrength$_{w20-24}$ [%]
I*
A
R = 0.812
ρ = 0.646
Legend
0 N
8 N
VEH
ML
BIS
PTH
SclAB
BIS + ML
PTH + ML
SclAB + ML

**Figure S.4: (Re)modeling velocity curves in mono- and combined treatments as an approximation of the mechanostat theory. (a)** (re)modeling velocity (RmV) curve fits (taking all animals per group into account) for VEH / ML, BIS / BIS + ML, PTH / PTH + ML, SclAB / SclAB + ML; **(b)** formation saturation level (FSL) **(c)** resorption saturation level (RSL), **(d)** (re)modeling threshold (RmT) **(b-d)** Each data point represents one animal (biological replicate; VEH: n=25, ML: n=26, BIS 0N: n=9, BIS 8N: n=9, PTH 0N: n=10, PTH 8N: n=9, SclAB 0N: n=9, SclAB 8N: n=8). The box spans the interquartile range (IQR) from the 25th to the 75th percentile. The median is shown as a line inside the box. Whiskers extend to the most extreme data points within 1.5 × IQR, points beyond this range are plotted as outliers. Δ denotes difference from VEH as no baseline value from week 20 was available. Bars show group means of sum of single vs. combined treatment indicating: antagonistic (I), additive (A) or synergistic (S) effects determined via treatment:loading interaction in a linear model on the absolute values. Significances were calculated using one-way ANOVA followed by Tukey’s post-hoc test; significant differences are depicted with a symbol in the color of the corresponding group (o: to monotreatment, ●: to combined treatment, see Supplementary Table S.1 for color encoding, p<0.05). **(g)** Linear regression of ΔStrength with FSL, and power-law regression with RmT in week interval w22-24. Source data are provided as a Source Data file.

**Table S.9:** Mechanostat parameters were derived from hyperbolic remodeling velocity curves as described by Marques et al.[3]. Formation (FSL) and resorption (RSL) saturation level curve fits were obtained by fitting each animal individually (see Statistical Methods section). One-way ANOVA in each time interval for the shown animal groups followed by Tukey’s post-hoc test was used on the absolute values. Synergy (S) or antagonism (I) between treatment and mechanical loading was assessed using the interaction term “treatment:loading” from a fitted linear model applied to the absolute values within each interval, because FSL and RSL can obtain negative values. ‘w20-22’ or ‘w22-24’ denotes the reference group; asterisks (*, **, ***) indicate groups that are significantly different from it. Significance is indicated as: *p<0.05, **p<0.01, ***p<0.001.

| Readout | N | w20-22 | w22-24 | %-change between w20-22 and w22-24 [%] | o | o | o | o | o | ● | ● | ● | Synergy w20-22 | Synergy w22-24 |
|---|---|---|---|---|---|---|---|---|---|---|---|---|---|---|
| **VEH (0N: n=25; 8N: n=26)** | | | | | | | | | | | | | | |
| FSL | 0N | -0.110 ± 0.030 | 0.089 ± 0.035 | -179.946 ± 55.587 | w20-22 | *** <0.001 | *** <0.001 | *** <0.001 | *** <0.001 | *** <0.001 | *** <0.001 | *** <0.001 | - | - |
| | | | | | w22-24 | *** <0.001 | 0.836 | *** <0.001 | *** <0.001 | *** <0.001 | *** <0.001 | *** <0.001 | | |
| | 8N | 0.268 ± 0.039 | 0.326 ± 0.030 | 39.760 ± 246.153 | *** <0.001 | w20-22 | 0.854 | 0.828 | 0.999 | 0.918 | *** <0.001 | *** <0.001 | - | - |
| | | | | | *** <0.001 | w22-24 | 0.067 | 0.826 | 0.138 | 0.968 | ** 0.003 | *** <0.001 | | |
| RSL | 0N | -1.564 ± 0.183 | -0.905 ± 0.143 | -13.991 ± 21.592 | w20-22 | 0.134 | *** <0.001 | 0.326 | 0.373 | *** <0.001 | *** <0.001 | * 0.010 | - | - |
| | | | | | w22-24 | 0.999 | ** 0.010 | *** <0.001 | * 0.03 | ** 0.003 | *** <0.001 | *** <0.001 | | |
| | 8N | -0.971 ± 0.200 | -0.810 ± 0.126 | 39.272 ± 32.069 | 0.134 | w20-22 | 0.096 | 1.000 | 1.000 | * 0.046 | *** <0.001 | 0.633 | - | - |
| | | | | | 0.999 | w22-24 | * 0.035 | *** <0.001 | 0.092 | * 0.012 | ** 0.002 | *** <0.001 | | |
| RmT | 0N | 593.596 ± 20.800 | 347.749 ± 39.075 | -37.136 ± 8.389 | w20-22 | 0.158 | *** <0.001 | ** 0.002 | * 0.011 | *** <0.001 | *** <0.001 | *** <0.001 | - | - |
| | | | | | w22-24 | 0.084 | *** <0.001 | *** <0.001 | *** <0.001 | *** <0.001 | *** <0.001 | *** <0.001 | | |
| | 8N | 441.163 ± 59.165 | 231.863 ± 33.763 | -32.738 ± 11.718 | 0.158 | w20-22 | 0.138 | 0.373 | 0.675 | *** <0.001 | * 0.039 | ** 0.003 | - | - |

| | | | | | | | | | | | | | | |
|---|---|---|---|---|---|---|---|---|---|---|---|---|---|---|
| | | | | | 0.084 | w22-24 | 0.112 | 0.068 | * 0.027 | * 0.029 | ** 0.009 | * 0.012 | | |
| **BIS (0N: n=9; 8N: n=9)** | | | | | | | | | | | | | | |
| FSL | 0N | 0.175 ± 0.051 | 0.166 ± 0.029 | -136.028 ± 75.652 | *** <0.001 | 0.854 | w20-22 | 1.000 | 0.997 | 0.380 | *** <0.001 | *** <0.001 | - | - |
| | | | | | 0.836 | 0.067 | w22-24 | ** 0.008 | *** <0.001 | * 0.04 | *** <0.001 | *** <0.001 | | |
| | 8N | 0.349 ± 0.050 | 0.374 ± 0.026 | 471.955 ± 478.813 | *** <0.001 | 0.918 | 0.380 | 0.347 | 0.810 | w20-22 | 0.056 | ** 0.005 | I* 0.033 | 0.715 |
| | | | | | *** <0.001 | 0.986 | * 0.04 | 1.000 | 0.818 | w22-24 | 0.176 | *** <0.001 | | |
| RSL | 0N | -0.109 ± 0.066 | -0.109 ± 0.075 | -82.421 ± 97.152 | *** <0.001 | 0.096 | w20-22 | 0.360 | 0.396 | 1.000 | 0.928 | 0.994 | - | - |
| | | | | | ** 0.010 | * 0.035 | w22-24 | 0.985 | 1.000 | 1.000 | 0.995 | 0.466 | | |
| | 8N | -0.023 ± 0.138 | -0.031 ± 0.048 | 111.987 ± 247.604 | *** <0.001 | * 0.096 | 0.360 | 0.396 | 1.000 | w20-22 | 0.928 | 0.994 | 0.245 | 0.945 |
| | | | | | ** 0.003 | * 0.012 | 1.000 | 0.999 | 0.999 | w22-24 | 1.000 | 0.657 | | |
| RmT | 0N | 225.688 ± 75.307 | 78.277 ± 29.808 | 44.450 ± 71.449 | *** <0.001 | 0.138 | w20-22 | 1.000 | 0.993 | 0.749 | 1.000 | 0.941 | - | - |
| | | | | | *** <0.001 | 0.112 | w22-24 | 1.000 | 1.000 | 1.000 | 0.996 | 0.995 | | |
| | 8N | 69.615 ± 26.501 | 50.264 ± 14.667 | 13.477 ± 26.845 | *** <0.001 | *** <0.001 | 0.749 | 0.397 | 0.254 | w20-22 | 0.932 | 1.000 | 0.971 | 0.246 |
| | | | | | *** <0.001 | * 0.029 | 1.000 | 1.000 | 1.000 | w22-24 | 1.000 | 1.000 | | |
| **PTH (0N: n=10; 8N: n=9)** | | | | | | | | | | | | | | |
| FSL | 0N | 0.176 ± 0.045 | 0.401 ± 0.029 | 123.157 ± 90.690 | *** <0.001 | 0.828 | 1.000 | w20-22 | 0.997 | 0.347 | *** <0.001 | *** <0.001 | - | - |
| | | | | | *** <0.001 | 0.826 | ** 0.008 | w22-24 | 0.959 | 1.000 | 0.354 | *** <0.001 | | |
| | 8N | 0.594 ± 0.062 | 0.542 ± 0.033 | -3.173 ± 8.406 | *** <0.001 | *** <0.001 | *** <0.001 | *** <0.001 | *** <0.001 | 0.056 | w20-22 | 0.985 | 0.641 | 0.207 |
| | | | | | *** <0.001 | ** 0.003 | *** <0.001 | 0.354 | 0.955 | 0.176 | w22-24 | 0.101 | | |
| RSL | 0N | -0.902 ± 0.189 | 0.125 ± 0.080 | -140.371 ± 26.161 | 0.326 | 1.000 | 0.360 | w20-22 | 1.000 | 0.233 | * 0.019 | 0.884 | - | - |
| | | | | | *** <0.001 | *** <0.001 | 0.985 | w22-24 | 0.927 | 0.999 | 1.000 | 0.928 | | |
| | 8N | 0.336 ± 0.192 | 0.089 ± 0.165 | 162.411 ± 219.057 | *** <0.001 | *** <0.001 | 0.928 | * 0.019 | * 0.025 | 0.977 | w20-22 | 0.519 | 0.131 | 0.656 |
| | | | | | *** <0.001 | ** 0.002 | 0.995 | 1.000 | 0.966 | 1.000 | w22-24 | 0.893 | | |
| RmT | 0N | 273.237 ± 43.684 | 73.302 ± 48.799 | -69.398 ± 20.353 | ** 0.002 | 0.373 | 1.000 | w20-22 | 1.000 | 0.397 | 0.984 | 0.704 | - | - |
| | | | | | *** <0.001 | 0.068 | 1.000 | w22-24 | 1.000 | 1.000 | 0.997 | 0.997 | | |
| | 8N | 186.112 ± 99.819 | 29.098 ± 5.321 | -32.602 ± 23.431 | *** <0.001 | * 0.039 | 1.000 | 0.984 | 0.929 | 0.932 | w20-22 | 0.994 | 0.564 | 0.330 |
| | | | | | *** <0.001 | ** 0.009 | 0.996 | 0.997 | 1.000 | 1.000 | w22-24 | 1.000 | | |
| **SclAB (0N: n=9; 8N: n=8)** | | | | | | | | | | | | | | |
| FSL | 0N | 0.230 ± 0.048 | 0.470 ± 0.043 | -141.051 ± 252.882 | *** <0.001 | 0.999 | 0.997 | 0.997 | w20-22 | 0.810 | *** <0.001 | *** <0.001 | - | - |
| | | | | | *** <0.001 | 0.138 | *** <0.001 | 0.959 | w22-24 | 0.818 | 0.955 | ** 0.004 | | |
| | 8N | 0.669 ± 0.071 | 0.732 ± 0.048 | 16.268 ± 10.997 | *** <0.001 | *** <0.001 | *** <0.001 | *** <0.001 | *** <0.001 | ** 0.005 | 0.985 | w20-22 | 0.508 | 0.734 |
| | | | | | *** <0.001 | *** <0.001 | *** <0.001 | *** <0.001 | ** 0.004 | *** <0.001 | 0.101 | w22-24 | | |

| | | | | | | | | | | | | | | |
|---|---|---|---|---|---|---|---|---|---|---|---|---|---|---|
| RSL | 0N | -0.900 ± 0.112 | -0.188 ± 0.208 | -62.023 ± 31.429 | 0.373 | 1.000 | 0.396 | 1.000 | w20-22 | 0.264 | * 0.025 | 0.898 | - | - |
| | | | | | * 0.03 | 0.092 | 1.000 | 0.927 | w22-24 | 0.999 | 0.966 | 0.293 | | |
| | 8N | -0.406 ± 0.296 | 0.446 ± 0.182 | -36.460 ± 82.623 | * 0.01 | 0.633 | 0.994 | 0.884 | 0.898 | 0.972 | 0.519 | w20-22 | 0.824 | 0.095 |
| | | | | | *** <0.001 | *** <0.001 | 0.466 | 0.928 | 0.293 | 0.657 | 0.893 | w22-24 | | |
| RmT | 0N | 303.569 ± 56.004 | 48.737 ± 18.175 | -84.419 ± 3.720 | * 0.011 | 0.675 | 0.993 | 1.000 | w20-22 | 0.254 | 0.929 | 0.530 | - | - |
| | | | | | *** <0.001 | * 0.027 | 1.000 | 1.000 | w22-24 | 1.000 | 1.000 | 1.000 | | |
| | 8N | 108.929 ± 28.828 | 25.493 ± 3.914 | -64.168 ± 8.188 | *** <0.001 | ** 0.003 | 0.941 | 0.704 | 0.530 | 1.000 | 0.994 | w20-22 | 0.716 | 0.233 |
| | | | | | *** <0.001 | * 0.012 | 0.995 | 0.997 | 1.000 | 1.000 | 1.000 | w22-24 | | |
